# Three-Dimensional Refractive Index Patterns Distinguish CD57-Defined Subsets of Human Memory T Cells

**DOI:** 10.64898/2026.11.16.767048

**Authors:** DongHun Ryu, Khoi Phuong Dao, Kyung Hwan Kim, Hayeon Cho, Young Seo Kim, Daewoong Ahn, Hyun-Seok Min, Geon Kim, Eui-Cheol Shin, YongKeun Park

## Abstract

Distinguishing T-cell subsets is important for characterizing immune differentiation, but label-free classification is challenging because closely related subsets have overlapping biophysical properties. We combined holotomography with a three-dimensional convolutional neural network to classify primary human T cells from their refractive index (RI) tomograms. Fluorescence-activated cell sorting defined four reference populations: CD4+, naïve CD8+, and CD57− and CD57+ memory CD8+ T cells. The FishNet-inspired classifier achieved 97.04% accuracy in a cell-level held-out test of 270 cells. Dedicated binary classifiers achieved 97.70% accuracy for naïve versus CCR7− memory CD8+ cells and 94.44% for CD57− versus CD57+ memory CD8+ cells. Although the memory subsets overlapped in whole-cell descriptors, their volumetric RI patterns supported classification. These findings establish a basis for investigating intrinsic optical phenotypes associated with CD57-defined T-cell differentiation.

**Abstract figure:** 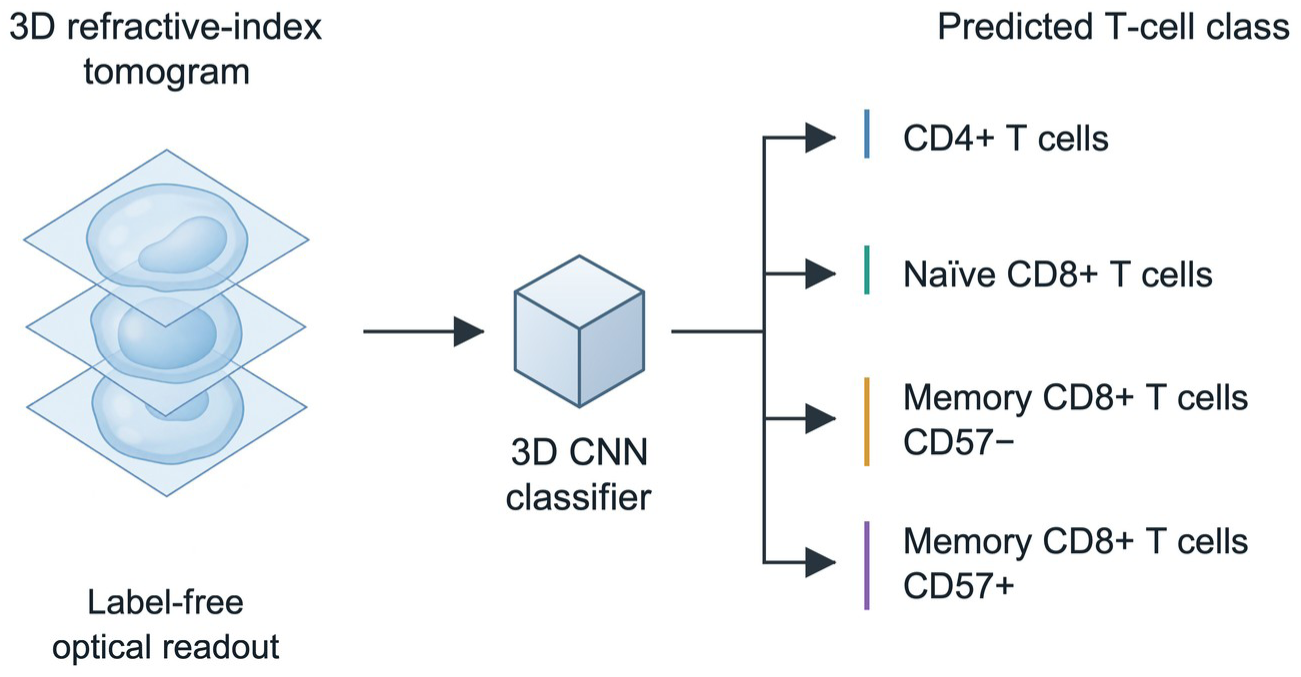

Conceptual illustration of three-dimensional (3-D) refractive index (RI) tomography and a convolutional neural network (CNN) for predicting four T-cell classes. The tomogram is schematic.

## 1 Introduction

T cells protect against pathogens by differentiating into populations with distinct immune functions [1]. Antigen exposure induces naïve T cells to proliferate and acquire effector functions, while persisting memory cells shape subsequent responses [2]. Repeated or chronic antigen exposure can lead to later differentiation states associated with CD57 expression [3, 4]. CD57 thus marks aspects of differentiation history, including cytomegalovirus (CMV) exposure, that relate to immune aging beyond chronological age [5]. Characterizing these states is relevant to studies of age-associated immune dysfunction and cancer immunotherapy [6], including chimeric antigen receptor (CAR)-T therapy [7]. Distinguishing CD57-defined subsets at the single-cell level could help characterize T-cell differentiation in these settings.

Flow cytometry identifies T-cell subsets by detecting cluster-of-differentiation (CD) markers with fluorescent antibodies [4, 8]. This molecular specificity provides reference identities for investigating other cellular properties. However, antibody engagement can alter live-cell function or phenotype, and staining, gating, and multiplexing require careful standardization [8, 9]. An optical readout based on intrinsic cellular properties could complement marker-based phenotyping by measuring cells without additional fluorescent contrast. The central challenge is whether such measurements capture enough information to distinguish closely related subsets.

Physical measurements can characterize cellular structure and composition, providing information beyond surface-marker identities. Raman spectroscopy detects molecular vibrations [10], while mass spectrometry identifies cell-surface glycoproteins [11]; acquisition time and specialized instrumentation can limit their use in cytometry. Quantitative phase imaging (QPI) measures the optical-path-length delay imposed by a cell [12], enabling biophysical studies of malaria-infected erythrocytes [13] and red-cell morphology in hematology [14]. Measurements of protein and lipid content by stimulated Raman scattering [15] and lipid quantification by QPI [16] further illustrate how optical contrast can report cellular composition. Machine learning has expanded the use of these measurements for cell and tissue classification and optical screening [17–23], while deep learning supports image reconstruction and restoration [24–28] and image analysis more broadly [29].

Holotomography (HT) provides a volumetric description of cellular optical properties. Whereas two-dimensional (2-D) QPI records projected phase delay, HT reconstructs three-dimensional (3-D) refractive index (RI) distributions from multiple optical measurements by solving an inverse scattering problem [30]. An RI volume therefore provides both whole-cell measurements and the spatial distribution of optical contrast within the cell. These complementary descriptions allow us to test whether spatial RI patterns distinguish T-cell subsets whose global descriptors overlap.

Label-free optical methods have distinguished T-cell populations that differ in lineage or activation state. Karandikar et al. distinguished naïve from anti-CD3/CD28-activated CD8+ T cells using diffraction phase microscopy [31], and Zhao et al. classified CD4+ and CD8+ T cells using cross-polarized diffraction imaging flow cytometry [32]. Deep-ultraviolet microscopy [33] and sub-millisecond holographic flow cytometry [34] have further expanded the available measurements. Here, we examine a finer distinction within the memory CD8+ population: whether volumetric RI patterns contain information associated with CD57-defined differentiation states.

We acquired RI tomograms from primary human T cells using HT, with fluorescence-activated cell sorting (FACS) defining four reference populations: CD4+, naïve CD8+, and CD57− and CD57+ memory CD8+ T cells (Fig. 1). We first characterized whole-cell morphology and biochemical descriptors, then trained a 3-D convolutional neural network (CNN) on the RI volumes. The network combines the multiscale architecture of FishNet [35], dense feature reuse from DenseNet [36], and channel attention from SENet [37] to integrate information across the cell volume.

**Figure 1.**
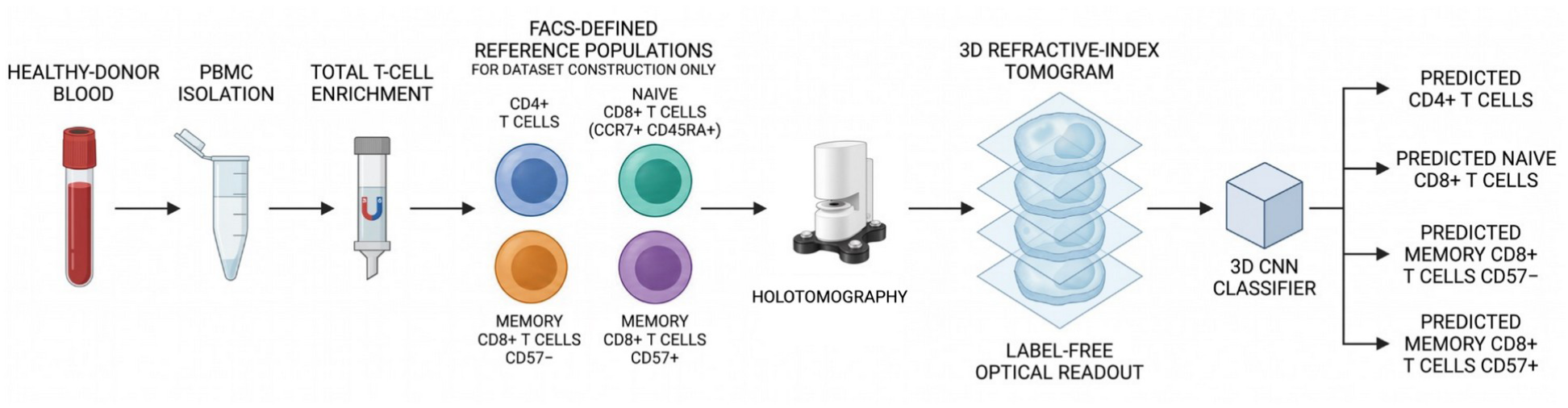
Workflow for label-free classification of human T-cell reference populations. Peripheral blood mononuclear cells (PBMCs) from healthy donors are isolated and enriched for T cells. Fluorescence-activated cell sorting (FACS) defines four reference populations: CD4+, naïve CD8+ (CCR7+CD45RA+), memory CD8+ CD57−, and memory CD8+ CD57+. Holotomography (HT) reconstructs three-dimensional (3-D) refractive index (RI) tomograms, which a 3-D convolutional neural network (CNN) uses to predict the reference class. FACS supplies the reference labels; the classifier receives RI volumes without fluorescence channels. Cell colors and the instrument drawing are schematic.

The four-class model achieved 97.04% accuracy in a cell-level held-out test. Dedicated binary models examined naïve versus memory CD8+ cells and the more closely related CD57− versus CD57+ memory subsets, testing discrimination at two levels of CD8+ differentiation. We also explored how subset fractions estimated by HT combined with artificial intelligence (HT + AI) compared with reported FACS fractions in mixed samples.

## 2 Materials and Methods

### Preparation of T-cell samples

Peripheral blood was obtained from healthy donors under approval from the Institutional Review Board (IRB) at the Korea Advanced Institute of Science and Technology (KAIST IRB-20-021), with informed consent from all participants. Peripheral blood mononuclear cells were isolated by Ficoll density-gradient centrifugation, and T cells were enriched by magnetic negative selection. Enriched cells were stained for CD3, CD4, CD8, CD45RA, CCR7, and CD57; dead cells were excluded using 7-aminoactinomycin D (7-AAD). Sorting on a FACSAria II instrument recovered CD4+ T cells, naïve (CCR7+CD45RA+) CD8+ T cells, and CD57− and CD57+ memory CD8+ T cells. Throughout this study, memory CD8+ denotes the sorted CCR7−CD45RA+/− fraction. Reagents and antibody clones are listed in Supplementary Methods S1. The mixed-sample measurements involved three donors, one per experimental day.

### Holotomography acquisition and reconstruction

RI tomograms were acquired with an HT-2H microscope (Tomocube, Inc., Republic of Korea), which implements coherent HT with a laser-based Mach–Zehnder interferometer (Supplementary Fig. 1). A 532 nm laser was split into sample and reference beams. A digital micromirror device controlled the sample illumination angle using digital grating patterns [38, 39]. Holograms were recorded at 49 illumination angles through matched water-immersion objectives with a numerical aperture of 1.2. The complete hologram sequence required approximately 0.1 s per cell. Optical components and the beam path are detailed in Supplementary Methods S2.

The complex optical field was recovered from each hologram [40, 41], and the 49 fields were combined by Fourier-diffraction-theorem inversion to reconstruct the 3-D RI distribution in TomoStudio [42]. Missing-cone artifacts were suppressed using 40 iterations of non-negativity regularization [43]. The lateral and axial resolutions were 110 nm and 355 nm, respectively [44, 45].

### Quantitative structural and biochemical parameter calculation

Single-cell structural and biochemical parameters were calculated from each 3-D RI tomogram. Surface area (SA) is denoted by S and volume by V. Fixed RI thresholding defined the cell mask; V was derived from the segmented interior and S from its boundary in the reconstructed spatial coordinates. These morphological descriptors complement the RI-derived biochemical measurements described below.

Sphericity (Ψ) was computed from S and V as

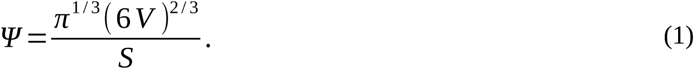

Biochemical descriptors were calculated using the linear relation between RI and the local concentration of nonaqueous molecules, predominantly proteins with contributions from lipids and nucleic acids. The protein density (PD) field *C* ( *r* ) was estimated from the voxel RI *n*( *r* ) and medium RI *n_m_*as

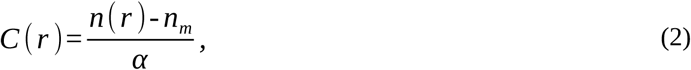

where α is the refractive index increment [46, 47]. We used *α* = 0.2 mL/g for protein-dominated cellular dry matter. Dry mass (DM) was obtained by integrating the PD over the segmented cellular volume (CV):

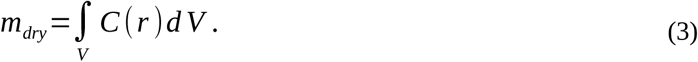

### Image processing and descriptive analysis

Image processing and feature extraction were performed in MATLAB R2018b, and RI isosurfaces were rendered in TomoStudio. Table 1 reports descriptor means and standard deviations; Fig. 2b shows descriptive distributions reconstructed from the original vector plots. The morphometry and learning cohorts and the provenance of the plotted observations are described in Supplementary Methods S4.

**Figure 2.**
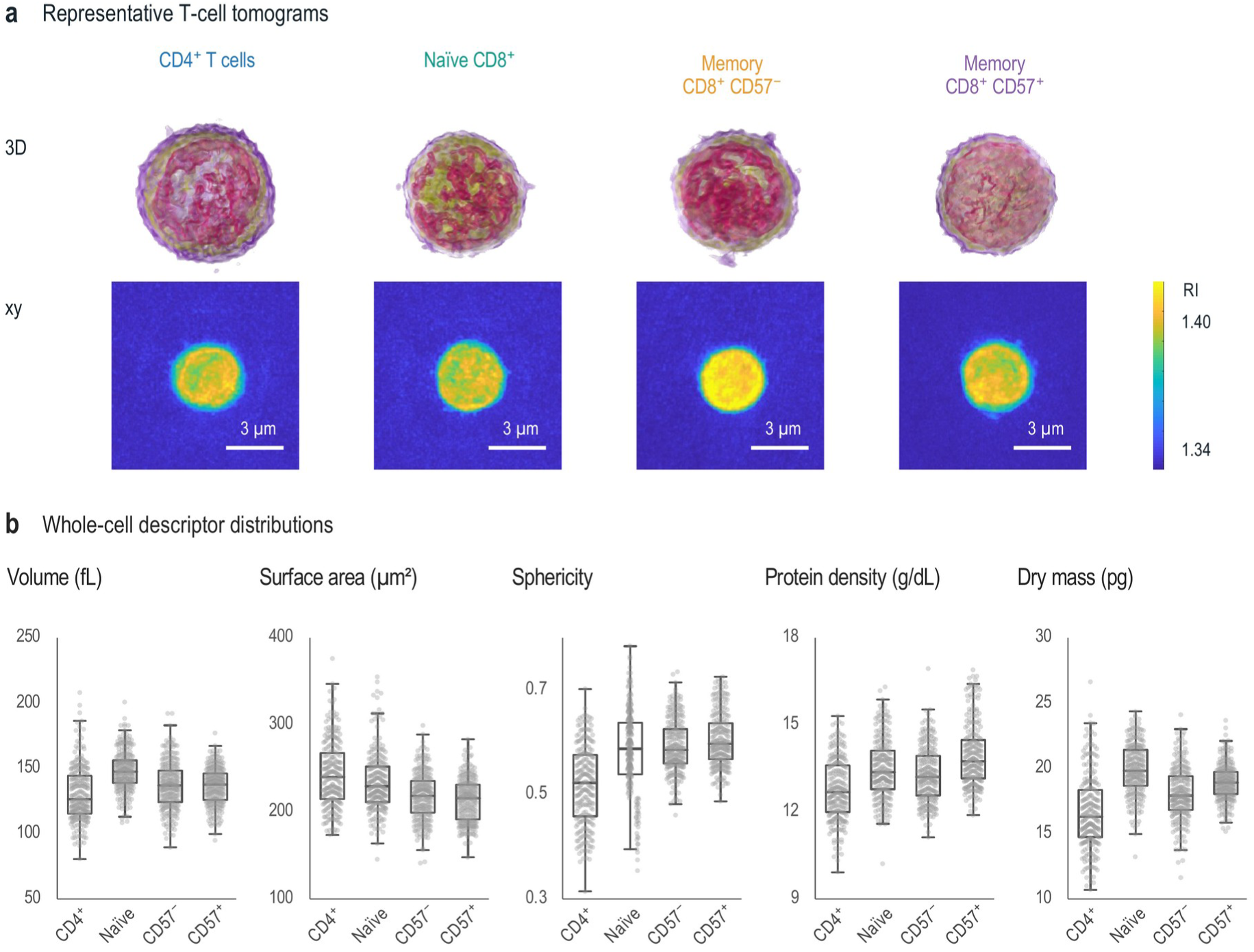
Representative refractive index tomograms and distributions of whole-cell descriptors. a, Representative three-dimensional (3-D) renderings (top) and xy refractive index (RI) sections (bottom) of CD4+, naïve CD8+, memory CD8+ CD57−, and memory CD8+ CD57+ T cells. The RI color scale spans 1.34–1.40. Scale bars, 3 µm. b, Distributions of cellular volume (CV, fL), surface area (SA, µmZ), sphericity, protein density (PD, g/dL), and dry mass (DM, pg), in the same class order. Boxes span the 25th–75th percentiles, center lines indicate medians, and whiskers extend to the most extreme observations within 1.5 times the interquartile range. Gray points represent individual observations, including outliers. CV, sphericity, PD, and DM contain 246 points per class; SA contains 245, 245, 246, and 246 points. Values were reconstructed from the original vector plots and are approximate. These plotted subsets differ from the cohort in Table 1 (Supplementary Methods S4).

**Table 1.**
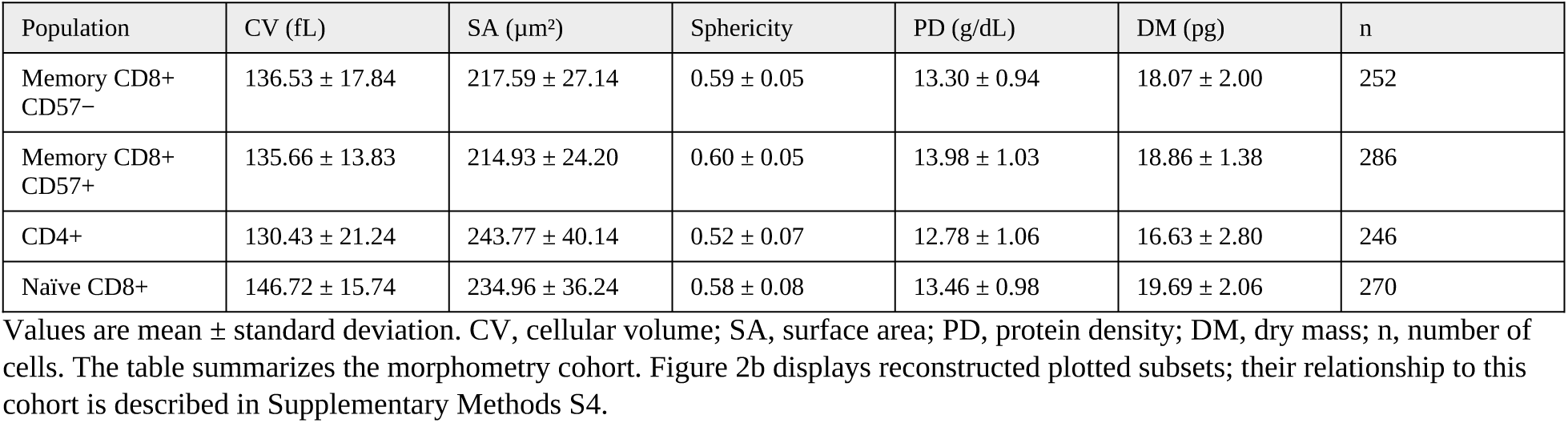
Cellular volume (CV), surface area (SA), sphericity, protein density (PD) and dry mass (DM) of the four T-cell populations.

### Conventional machine-learning baselines

We evaluated k-nearest neighbor (k-NN) and support vector machine (SVM) classifiers using features derived from CV, SA, sphericity, PD, and DM. For k-NN, evaluating the five descriptors at 20 RI thresholds produced a 100-dimensional feature vector; the reported classifier used k = 5 (Supplementary Fig. 2). Binary and multiclass SVMs were implemented with MATLAB’s fitcsvm and fitcecoc functions, respectively. Kernel type, box constraint, and kernel scale were tuned separately for each task by cross-validation on the training partition. Feature-extraction settings and test counts are provided in Supplementary Methods S3 and Supplementary Table 3.

The conventional classifiers and CNNs used different model-specific test partitions. We therefore report their accuracies as descriptive comparisons across the same classification tasks, rather than paired comparisons on identical test cells.

### 3-D deep-learning classifier

The CNN processes RI volumes through a FishNet-inspired tail–body–head architecture [35] (Fig. 3a). The tail downsamples the input and extracts features with DenseNet-style connectivity [36]. The body upsamples these features and combines them with higher-resolution representations through cross-stage connections and residual convolutional blocks. The head then downsamples and refines features from both earlier stages. SENet-style channel attention weights features at the tail–body bridge [37] (Fig. 3b). A decision block combines batch normalization, LeakyReLU activation, convolution, and adaptive average pooling; a final convolution and softmax yield class probabilities (Fig. 3c). This design allows the classifier to combine information at multiple spatial scales. The layer specification is given in Supplementary Table 1.

**Figure 3.**
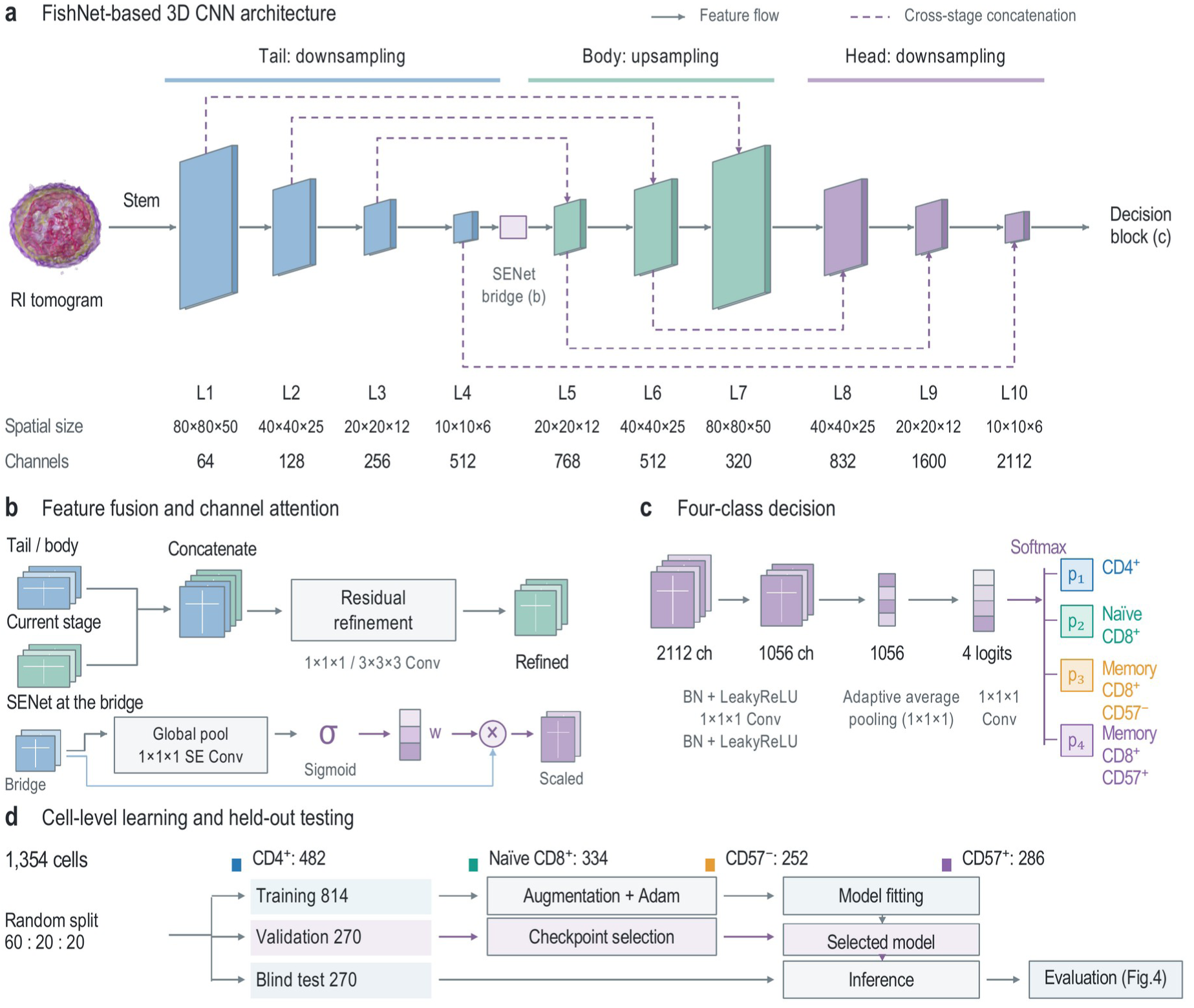
FishNet-based architecture and cell-level evaluation workflow. a, A refractive index (RI) tomogram passes through a convolutional stem, downsampling tail (L1–L4), upsampling body (L5–L7), downsampling head (L8–L10), and decision block. Solid arrows indicate sequential feature flow; dashed arrows indicate cross-stage channel concatenation between L1–L7, L2–L6, L3–L5, L6–L8, L5–L9, and L4–L10. Spatial dimensions and channel counts are given below each level; feature planes are schematic. b, Cross-stage features are concatenated and refined by residual convolutional blocks. At the L4–L5 bridge, squeeze-and-excitation (SENet) attention uses global pooling, 1 × 1 × 1 convolutions, and a sigmoid to generate channel weights that multiply the bridge features. c, The decision block projects 2,112 channels to 1,056 using batch normalization (BN), LeakyReLU activation, and a 1 × 1 × 1 convolution. Adaptive average pooling and a final convolution produce four logits, which softmax converts into class probabilities. Weights and probabilities are schematic. d, The four-class learning cohort comprises 1,354 cells: 482 CD4+, 334 naïve CD8+, 252 memory CD8+ CD57−, and 286 memory CD8+ CD57+. The cell-level split contains 814 training and 270 held-out-test cells; 270 validation cells are inferred from the total. Validation selects the checkpoint for testing. Dedicated binary models use separate task-specific partitions (Supplementary Table 3).

The four-class dataset contained 1,354 tomograms and was randomly partitioned at the cell level according to a 6:2:2 training/validation/held-out-test scheme. The records specify 814 training and 270 test cells, leaving 270 for validation (Fig. 3d). Training used the adaptive moment estimation (Adam) optimizer, and validation accuracy selected the checkpoint for held-out testing. Augmentation included random translation, cropping, elastic transformation, and additive Gaussian noise. The implementation used PyTorch 1.0.1. Dedicated binary models used task-specific partitions, with 87 test cells for naïve versus memory CD8+ and 54 for the CD57 contrast. Training settings and the available partition counts are reported in Supplementary Methods S3 and Supplementary Table 3. The evaluation used cell-level separation; donor membership across partitions was not documented.

### Exploratory comparison of mixed-sample fractions

We compared FACS and HT + AI subset fractions from three donors measured on separate experimental days (Fig. 5). The mixed-sample model incorporated additional unsorted Day 1 cells assigned an assumed CD4+ label because the corresponding FACS sample was CD4+-enriched. Thus, Day 1 contributed to model development. The reported fractions were plotted without renormalization; the development history and available fraction records are detailed in Supplementary Methods S5 and Supplementary Table 5.

### Visualization of network outputs

For a qualitative view of network outputs, we randomly sampled 120 tomograms, 30 per reference class, and embedded their four-component final-output vectors in two dimensions using t-distributed stochastic neighbor embedding (t-SNE) in MATLAB R2018b (Fig. 4d). The plotted symbols identify the FACS reference classes.

**Figure 4.**
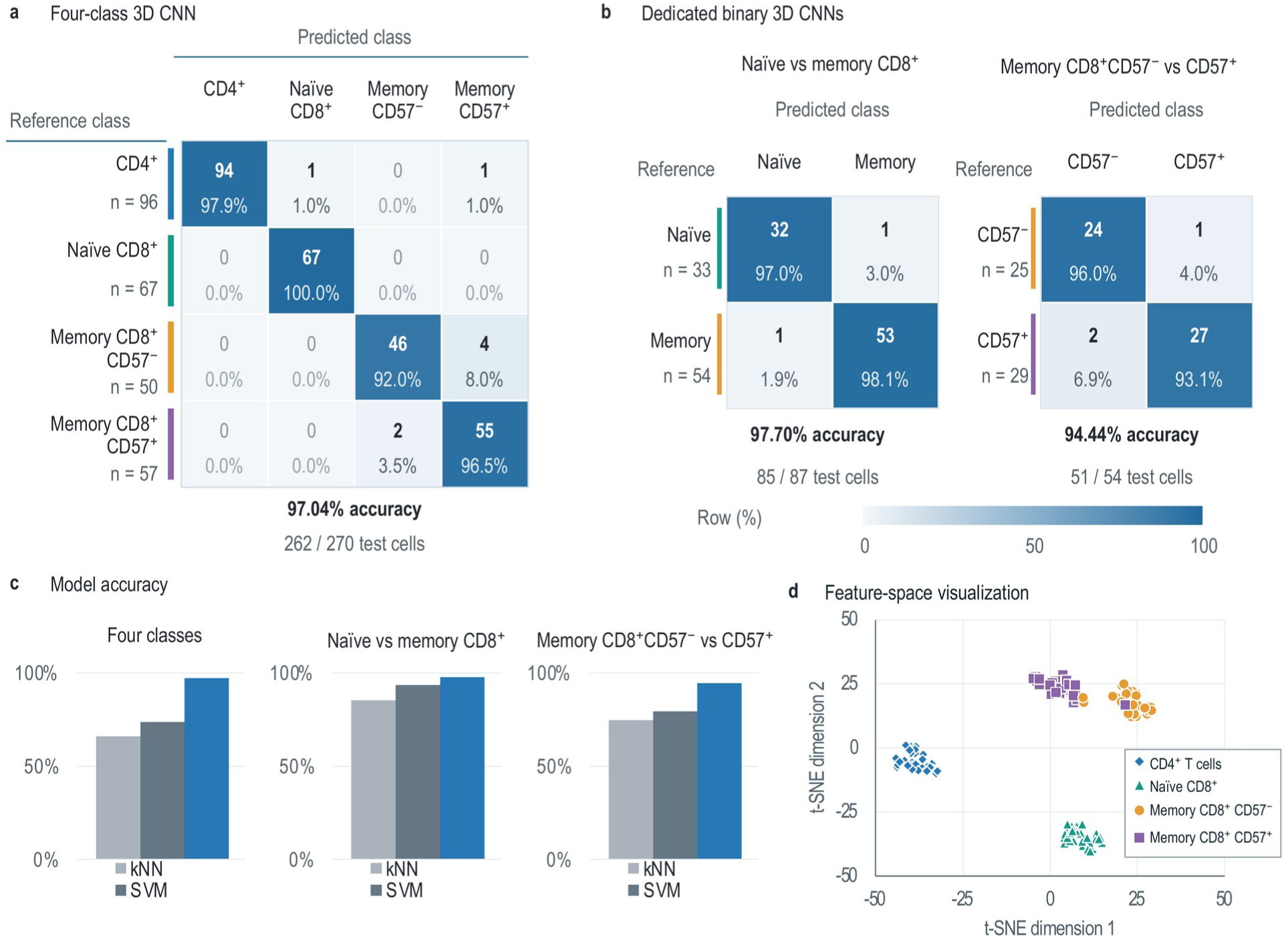
Classification performance and network-output visualization. a, Four-class convolutional neural network (CNN) confusion matrix for 270 blind-test cells. Rows denote fluorescence-activated cell sorting (FACS) reference classes and columns denote predictions, ordered CD4+, naïve CD8+, memory CD8+ CD57− and memory CD8+ CD57+; row sample sizes are 96, 67, 50 and 57. Entries show counts and percentages normalized within each reference row. Overall accuracy is 97.04% (262/270); six of eight errors occur between the two memory CD8+ subsets. b, Confusion matrices for dedicated binary CNNs: naïve versus memory CD8+ T cells, 97.70% (85/87; reference row sizes 33 and 54), and memory CD8+ CD57− versus CD57+ T cells, 94.44% (51/54; row sizes 25 and 29). All matrices share a 0–100% row-normalized color scale. c, Test accuracies for k-nearest neighbors (k-NN), support vector machines (SVM), and the three-dimensional (3-D) CNN, ordered left to right within each task. Correct/test-cell counts and exact accuracies are listed in Supplementary Table 3. Model-specific test partitions and sample sizes differ, so the comparison is descriptive. d, Two-dimensional (2-D) t-distributed stochastic neighbor embedding (t-SNE) of four-component final network outputs for 120 cells, 30 from each FACS-defined reference population. Blue diamonds, CD4+; teal triangles, naïve CD8+; orange circles, memory CD8+ CD57−; purple squares, memory CD8+ CD57+. Symbols encode reference identity. Axes are arbitrary embedding coordinates.

**Figure 5.**
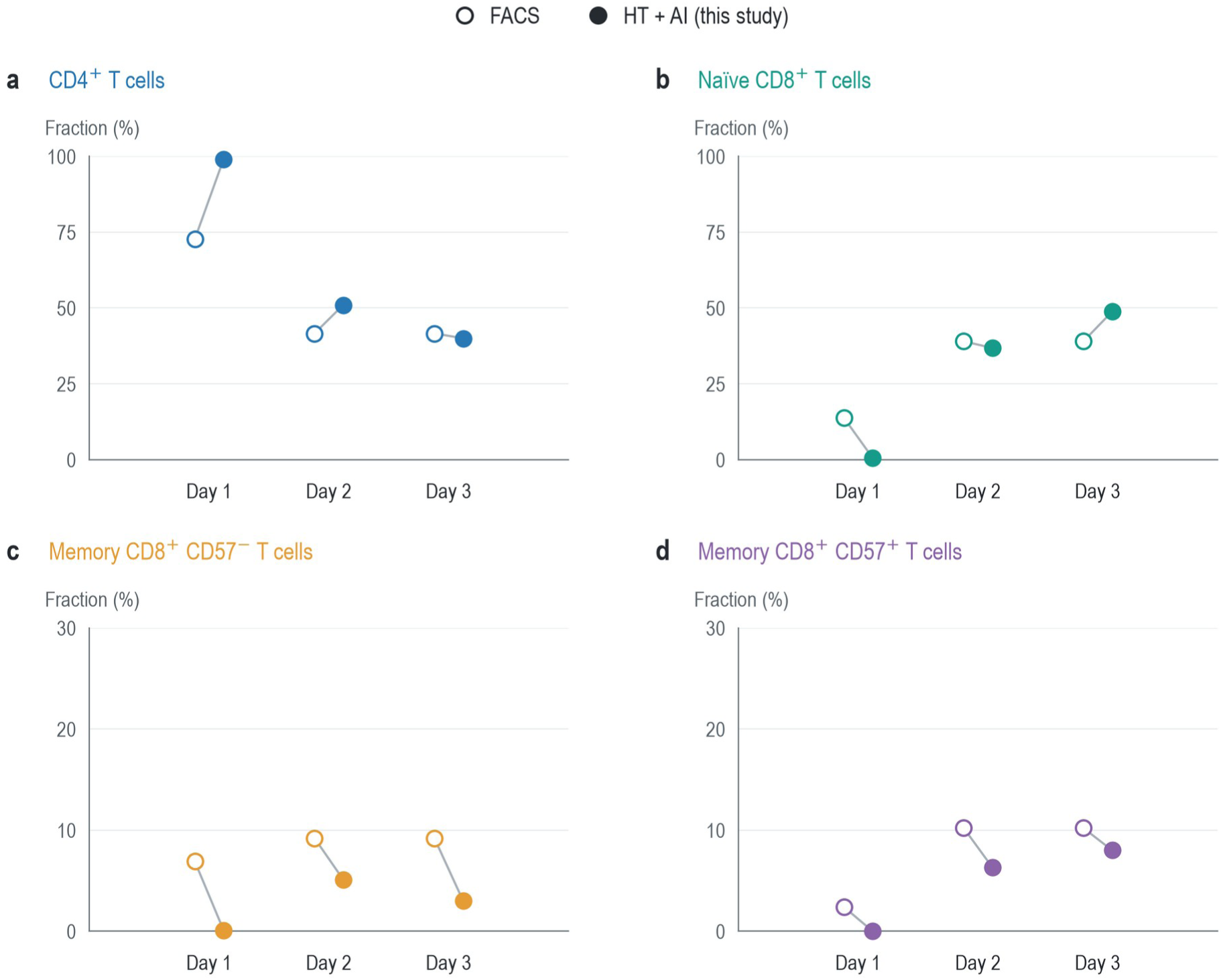
Exploratory comparison of reported FACS and HT + AI T-cell subset fractions from three donors. a, CD4+ T cells. b, Naïve CD8+ T cells. c, Memory CD8+ CD57− T cells. d, Memory CD8+ CD57+ T cells. Fluorescence-activated cell sorting (FACS) provides the reported comparison fractions. Holotomography (HT) combined with artificial intelligence (AI) classification is labeled HT + AI. Open circles denote FACS and filled circles denote HT + AI (this study). Gray segments connect the two estimates within each experimental day. Horizontal offsets separate the methods and do not encode a measured variable. Panels a and b use 0–100% axes, whereas panels c and d use 0–30% axes. Fractions are shown without renormalization and may sum to less than 100%. Exact values are listed in Supplementary Table 5. Days 1–3 represent three donors, one per day; each point is a donor-level fraction. Unsorted Day 1 cells contributed to model development with assumed CD4+ labels. Development–evaluation overlap and the methods’ fraction denominators and inclusion criteria could not be verified. The comparison is exploratory (Supplementary Methods S5).

## 3 Results

The four FACS-defined populations span a CD4+/CD8+ distinction and two comparisons within CD8+ cells: naïve versus memory and CD57− versus CD57+ memory. We used these comparisons to examine how classification changes as the phenotypes become more closely related. Representative RI volumes and xy sections illustrate the optical measurements available for each population (Fig. 2a). We first analyzed whole-cell descriptors and their classification performance, then evaluated the information retained in the full RI volumes.

### Whole-cell descriptors overlap across memory T-cell subsets

To characterize morphology and composition across the four populations, we quantified CV, SA, sphericity, PD, and DM from the RI tomograms. Population-level differences in these descriptors coexisted with substantial overlap among individual cells. Table 1 summarizes 246 CD4+, 270 naïve CD8+, 252 CD57− memory CD8+, and 286 CD57+ memory CD8+ cells. Figure 2b displays the available plotted subsets, whose counts differ from the Table 1 cohort (Supplementary Methods S4).

CD4+ cells showed a smaller mean volume with a larger mean surface area than the CD8+ populations. Their mean CV was 130.43 ± 21.24 fL, compared with 146.72 ± 15.74 fL for naïve CD8+ cells, while their mean SA was 243.77 ± 40.14 µmZ, compared with 214.93 ± 24.20 µmZ for CD57+ memory CD8+ cells. This relationship between surface area and volume is reflected in the lower sphericity of CD4+ cells, approximately 0.52 versus 0.58–0.60 for the CD8+ populations. CD4+ cells also had lower mean PD and DM (Table 1). Thus, the RI measurements captured differences in both overall cell shape and RI-derived composition across the CD4+/CD8+ distinction.

Within the CD8+ populations, the CD57− and CD57+ memory subsets had similar mean volumes, surface areas, and sphericities. Their mean volumes were 136.53 and 135.66 fL, respectively. CD57+ memory cells had a higher mean PD than CD57− cells, but the distributions overlapped. Naïve CD8+ and CD57− memory cells also showed similar sphericity and PD. These observations explain why differences in population means need not yield reliable classification of individual cells: a cell’s descriptor values can be compatible with more than one subset.

To examine whether combinations of these descriptors could resolve the overlapping populations, we evaluated k-NN and SVM classifiers (Fig. 4c; Supplementary Figs 2–5). Four-class accuracies were 66.19% and 73.81%, respectively. Both methods performed better on naïve versus memory CD8+ cells, reaching 85.19% and 93.52%, but their accuracies fell to 74.77% and 79.44% for the CD57 contrast. The lower performance on the latter task is consistent with the close similarity of the memory subsets in the descriptor distributions. These baselines establish the difficulty of distinguishing CD57-defined populations using the selected whole-cell features and motivate analysis of the full RI volumes. Correct/test-cell counts are listed in Supplementary Table 3.

### Volumetric classification distinguishes the reference populations

To test whether spatial RI patterns could distinguish populations with overlapping whole-cell descriptors, we trained a volumetric CNN. The network combines multiscale processing from FishNet [35], dense feature reuse [36], and channel attention [37] to integrate RI patterns across the cell (Fig. 3). We evaluated a four-class model and dedicated models for the two CD8+ comparisons. The four-class learning cohort contained 1,354 cells; validation selected the checkpoint before evaluation on 270 held-out cells. The binary models were trained and evaluated with their own task-specific partitions (Methods; Supplementary Table 3).

The four-class CNN correctly classified 262 of 270 test cells, yielding 97.04% accuracy (Fig. 4a). All 67 naïve CD8+ cells and 94 of 96 CD4+ cells were correctly identified. Most residual errors occurred within the memory CD8+ population: four CD57− cells were predicted as CD57+, and two CD57+ cells were predicted as CD57−. These exchanges accounted for six of the eight errors, showing that the closest phenotypic comparison remained the principal source of confusion even at high overall accuracy. Class-specific recalls and training performance are reported in Supplementary Table 4.

In the t-SNE visualization, final network outputs formed largely separated groups corresponding to the four reference populations (Fig. 4d). The two memory CD8+ groups were adjacent, and two CD57− reference cells lay near the CD57+ group, illustrating the proximity of some outputs across this boundary. The symbols denote reference identity and the embedding represents output similarity; prediction errors are quantified separately in the confusion matrix.

Dedicated binary classifiers examined the two CD8+ distinctions directly (Fig. 4b). Accuracy was 97.70% for naïve versus memory cells (85/87) and 94.44% for CD57− versus CD57+ memory cells (51/54). The lower accuracy on the CD57 task agrees with the concentration of four-class errors between the memory subsets. Nevertheless, most cells in this closely related pair were correctly distinguished from their RI volumes despite the overlapping global descriptors. The CNN accuracies were higher than the corresponding k-NN and SVM summaries (Fig. 4c); for the CD57 contrast, the CNN and SVM values differed by 15.00 percentage points. Model-specific test partitions differed, so this difference describes the reported results across models rather than a paired estimate of improvement on the same cells.

A DenseNet classifier [36] provided an additional deep-learning baseline, with reported four-class test accuracies of 78–83% (Supplementary Table 2), below the FishNet-inspired model’s accuracy. The recovered summary does not specify the settings and partition membership associated with this range. A matched evaluation with controlled ablations would be needed to attribute the performance difference to multiscale processing, dense connections, or channel attention.

### Exploratory comparison of reported T-cell subset fractions

To examine whether HT + AI classifications reproduced FACS subset fractions in mixed samples, we compared reported fractions from three donors (Fig. 5). The largest discrepancy occurred on Day 1: HT + AI assigned 99.0% of cells to CD4+, whereas FACS reported 72.7%, a difference of 26.3 percentage points. The corresponding naïve CD8+ fractions were 0.6% and 13.8%. On Days 2 and 3, the reported distributions included substantial CD4+ and naïve CD8+ fractions with both methods, although their relative proportions differed. The full set of day-specific values is provided in Supplementary Table 5.

Both memory subsets had lower HT + AI fractions than FACS fractions on all three days. The CD57− differences ranged from −6.8 to −4.1 percentage points, and the CD57+ differences from −3.9 to −2.2 percentage points. These consistent directions of disagreement show that high accuracy on sorted reference cells did not by itself establish agreement in mixed-sample composition. Day 1 cells had contributed to model development with assumed CD4+ labels, and the methods’ fraction denominators and inclusion criteria could not be verified. Figure 5 therefore identifies discrepancies for further investigation; it does not provide an independent estimate of composition accuracy.

## 4 Discussion

Volumetric RI patterns distinguished CD57-defined memory CD8+ populations whose whole-cell descriptors overlapped. The four-class analysis showed separation across a broader set of T-cell populations, while the dedicated CD57 classifier tested the distinction within memory CD8+ cells directly. Its 94.44% test accuracy indicates that the measured volumes contain an intrinsic optical phenotype associated with the FACS-defined CD57 classes. The concentration of four-class errors between the memory subsets is consistent with their more similar optical representations and identifies this finer distinction as the principal source of confusion.

Spatial RI organization may explain why volumetric classification distinguished subsets with overlapping global descriptors. Volume, surface area, sphericity, mean protein density, and dry mass summarize the cell with a small number of values; even their evaluation at multiple RI thresholds does not retain the full spatial arrangement of the contrast. The CNN can combine local RI patterns across spatial scales. Cells with similar overall size and dry mass could consequently produce different network outputs if their intracellular RI distributions differ. Identifying which spatial features contribute to the CD57 distinction would test this interpretation directly.

The classifier used RI data without fluorescence inputs, but the imaged cells had already undergone antibody staining and FACS sorting. Because labeling and preparation can affect cellular properties [8, 9], evaluation on cells measured before staining would clarify how well the learned phenotype transfers to an entirely label-free workflow. Donor-held-out studies and balanced preparation and acquisition conditions would further separate subset-associated features from donor or batch effects.

The CD57 result extends optical T-cell phenotyping to closely related memory CD8+ subsets with limited separation in global morphology. Diffraction phase microscopy distinguished naïve from activated CD8+ cells [31], and diffraction imaging flow cytometry distinguished CD4+ from CD8+ cells [32]. Holographic flow cytometry [34] and deep-ultraviolet microscopy [33] broadened the speed and contrast mechanisms available for T-cell measurements. Related 3-D RI studies addressed leukocyte identity [48], CAR-T immunological synapses [49], and CD8+ morphology in sepsis [50]. CD57’s association with terminal differentiation and CMV exposure [5] gives this contrast biological relevance, while its links to immune aging and cancer [6] motivate examining how the optical phenotype relates to functional state. Such interpretation requires measurements of differentiation and function beyond CD57 expression alone.

Donor-independent evaluation with CMV serology would test whether the optical distinction persists across different immune histories. CMV serostatus was not controlled in this healthy-donor cohort, although it is associated with CD57 expression [5]. At the cellular level, feature-attribution methods [51–53] could identify regions associated with the classifier’s decisions. Controlled perturbations of the RI input and matched model comparisons could then test how those regions contribute to classification.

Practical use also depends on the full measurement pipeline. Acquisition of 49 holograms took approximately 0.1 s per cell, but throughput includes cell handling, reconstruction, preprocessing, and inference. Integrating these stages with flow-based delivery or faster detection could support larger-scale phenotyping. Evaluation should therefore measure both end-to-end performance and the suitability of the imaged cells for subsequent functional assays.

Population-level measurements require subset fractions that remain calibrated as sample composition and cell preparation change. The Day 1 CD4+ discrepancy and the lower estimated memory fractions across all three days identify this as an unresolved step beyond single-cell discrimination. Because Day 1 contributed to development and the fraction denominators were unavailable, the current comparison cannot determine whether the differences arose from classification, sampling, or inclusion criteria. A prospective comparison using independent donors, predefined inclusion criteria, and matched aliquots would directly test quantitative agreement between HT-based estimates and FACS.

## 5 Conclusion

Holotomography combined with a 3-D CNN distinguished four human T-cell populations in cell-level held-out tests, including CD57− and CD57+ memory CD8+ cells with overlapping whole-cell descriptors. The findings support volumetric RI patterns as a source of information for T-cell phenotyping and motivate studies linking these patterns to differentiation and function across independent donors.

## Data and code availability

The data that support the findings of this study are available from the corresponding author upon request. The training and evaluation code and trained model weights are available from the corresponding author upon request. Reuse is subject to the license terms of the underlying software. The implementation used PyTorch 1.0.1 and MATLAB R2018b.

## AI assistance

OpenAI Codex was used to assist with language editing. The authors retain responsibility for verifying the final text, citations and scientific interpretation.

## Acknowledgments

This work was supported by KAIST, Tomocube, Inc., the National Research Foundation of Korea (NRF) grant funded by the Korean government (Ministry of Science and ICT; RS-2024-00442348), and the Ministry of Trade, Industry and Energy and Korea Institute for Advancement of Technology through the International Cooperative R&D program (P0028463). Y.K.P. acknowledges support from a Merck fellowship. The authors thank members of the Park and Shin laboratories for helpful discussions and the KAIST Analysis Center for Research Advancement for technical support.

## Author contributions

E.-C.S. and Y.K.P. conceived the study and supervised the project. D.H.R., K.H.K., K.P.D., and H.C. designed and performed the experiments. K.H.K. prepared and FACS-purified the primary human T-cell populations under E.-C.S.’s supervision. D.H.R. and Y.S.K. performed the 3-D RI tomographic imaging and reconstruction. K.P.D. developed and implemented the classical-machine-learning baselines (k-NN and SVM). H.C. developed the 3-D convolutional neural network and performed the deep-learning experiments. D.A. and H.-S.M. contributed to the design and analysis of the imaging and classification pipelines. G.K. refined the classifier implementation and regenerated the figure panels used in the current version of the manuscript. E.-C.S. provided the immunological interpretation of the results. D.H.R., K.H.K., K.P.D., E.-C.S., and Y.K.P. wrote the manuscript.

## Competing interests

K.P.D., Y.S.K., D.A., H.-S.M., G.K., and Y.K.P. have financial interests in Tomocube, Inc., a company that commercializes holotomography and quantitative phase imaging instruments and that is one of the sponsors of this work. Y.K.P. is a co-founder, a shareholder, and currently serves in a scientific-advisory role at Tomocube, Inc. D.A. and H.-S.M. are employees of Tomocube, Inc. K.P.D., Y.S.K., and G.K. hold Tomocube-related equity as former or current affiliates. The remaining authors (D.H.R., K.H.K., H.C., and E.-C.S.) declare no competing interests.

## Supporting Information

- Supplementary Methods S1–S5: reagents, optical configuration, model training, cohort definitions, and mixed-sample records.
- Supplementary Figures 1–5: coherent holotomography optical system, k-nearest neighbor workflow and results, and support vector machine results.
- Supplementary Tables 1–2: network layer specification and reported classifier accuracies.
- Supplementary Tables 3–4: test counts, task-specific partitions, training performance, and class-specific recalls.
- Supplementary Table 5: FACS and HT + AI subset fractions by experimental day.

## Table of contents entry

Three-Dimensional Refractive Index Patterns Distinguish CD57-Defined Subsets of Human Memory T Cells

DongHun Ryu+, Khoi Phuong Dao+, Kyung Hwan Kim, Hayeon Cho, Young Seo Kim, Daewoong Ahn, Hyun-Seok Min, Geon Kim, Eui-Cheol Shin and YongKeun Park*

Holotomography and a three-dimensional convolutional neural network classify four human T-cell populations from refractive index tomograms, achieving 97.04% accuracy in a cell-level held-out test. Volumetric patterns distinguish CD57− and CD57+ memory CD8+ subsets despite overlapping whole-cell descriptors, linking intrinsic optical measurements to marker-defined phenotypes.

## References

[1] J. E. Smith-Garvin, G. A. Koretzky, and M. S. Jordan, T Cell Activation, Annu. Rev. Immunol., 2009, 27, 591–619, doi:10.1146/annurev.immunol.021908.132706.

[2] S. M. Kaech, and W. Cui, Transcriptional control of effector and memory CD8+ T cell differentiation, Nat. Rev. Immunol., 2012, 12, 749–761, doi:10.1038/nri3307.

[3] A. N. Akbar, S. M. Henson, and A. Lanna, Senescence of T Lymphocytes: Implications for Enhancing Human Immunity, Trends Immunol., 2016, 37, 866–876, doi:10.1016/j.it.2016.09.002.

[4] N. Weng, A. N. Akbar, and J. Goronzy, CD28− T cells: their role in the age-associated decline of immune function, Trends Immunol., 2009, 30, 306–312, doi:10.1016/j.it.2009.03.013.

[5] G. Autaa, D. Korenkov, J. van Beek, I. Pellegrin, B. Parfait, D. van Baarle, O. Launay, E. Tartour, and V. Appay, Re-evaluating CD57 as a marker of T cell senescence: implications for immune ageing and differentiation, Immun. Ageing, 2025, 22, 47, doi:10.1186/s12979-025-00542-8.

[6] Q. Liu, J. Li, X. Sun, J. Lin, Z. Yu, Y. Xiao, D. Li, B. Sun, H. Bao, and Y. Liu, Immunosenescence and cancer: molecular hallmarks, tumor microenvironment remodeling, and age-specific immunotherapy challenges, J. Hematol. Oncol., 2025, 18, 81, doi:10.1186/s13045-025-01735-w.

[7] C. H. June, R. S. O’Connor, O. U. Kawalekar, S. Ghassemi, and M. C. Milone, CAR T cell immunotherapy for human cancer, Science, 2018, 359, 1361–1365, doi:10.1126/science.aar6711.

[8] A. Cossarizza, H. Chang, A. Radbruch, M. Akdis, I. Andrä, F. Annunziato, P. Bacher, V. Barnaba, L. Battistini, W. M. Bauer, S. Baumgart, B. Becher, W. Beisker, C. Berek, A. Blanco, G. Borsellino, P. E. Boulais, R. R. Brinkman, M. Büscher, D. H. Busch, T. P. Bushnell, X. Cao, A. Cavani, P. K. Chattopadhyay, Q. Cheng, S. Chow, M. Clerici, A. Cooke, A. Cosma, L. Cosmi, A. Cumano, V. D. Dang, D. Davies, S. De Biasi, G. Del Zotto, S. Della Bella, P. Dellabona, G. Deniz, M. Dessing, A. Diefenbach, J. Di Santo, F. Dieli, A. Dolf, V. S. Donnenberg, T. Dörner, G. R. A. Ehrhardt, E. Endl, P. Engel, B. Engelhardt, C. Esser, B. Everts, A. Dreher, C. S. Falk, T. A. Fehniger, A. Filby, S. Fillatreau, M. Follo, I. Förster, J. Foster, G. A. Foulds, P. S. Frenette, D. Galbraith, N. Garbi, M. D. García-Godoy, J. Geginat, K. Ghoreschi, L. Gibellini, C. Goettlinger, C. S. Goodyear, A. Gori, J. Grogan, M. Gross, A. Grützkau, D. Grummitt, J. Hahn, Q. Hammer, A. E. Hauser, D. L. Haviland, D. Hedley, G. Herrera, M. Herrmann, F. Hiepe, T. Holland, P. Hombrink, J. P. Houston, B. F. Hoyer, B. Huang, C. A. Hunter, A. Iannone, H. Jäck, B. Jávega, S. Jonjic, K. Juelke, S. Jung, T. Kaiser, T. Kalina, B. Keller, S. Khan, D. Kienhöfer, T. Kroneis, D. Kunkel, C. Kurts, P. Kvistborg, J. Lannigan, O. Lantz, A. Larbi, S. LeibundGut-Landmann, M. D. Leipold, M. K. Levings, V. Litwin, Y. Liu, M. Lohoff, G. Lombardi, L. Lopez, A. Lovett-Racke, E. Lubberts, B. Ludewig, E. Lugli, H. T. Maecker, G. Martrus, G. Matarese, C. Maueröder, M. McGrath, I. McInnes, H. E. Mei, F. Melchers, S. Melzer, D. Mielenz, K. Mills, D. Mirrer, J. Mjösberg, J. Moore, B. Moran, A. Moretta, L. Moretta, T. R. Mosmann, S. Müller, W. Müller, C. Münz, G. Multhoff, L. E. Munoz, K. M. Murphy, T. Nakayama, M. Nasi, C. Neudörfl, J. Nolan, S. Nourshargh, J. O’Connor, W. Ouyang, A. Oxenius, R. Palankar, I. Panse, P. Peterson, C. Peth, J. Petriz, D. Philips, W. Pickl, S. Piconese, M. Pinti, A. G. Pockley, M. J. Podolska, C. Pucillo, S. A. Quataert, T. R. D. J. Radstake, B. Rajwa, J. A. Rebhahn, D. Recktenwald, E. B. M. Remmerswaal, K. Rezvani, L. G. Rico, J. P. Robinson, C. Romagnani, A. Rubartelli, B. Ruckert, J. Ruland, S. Sakaguchi, F. Sala-de-Oyanguren, Y. Samstag, S. Sanderson, B. Sawitzki, A. Scheffold, M. Schiemann, F. Schildberg, E. Schimisky, S. A. Schmid, S. Schmitt, K. Schober, T. Schüler, A. R. Schulz, T. Schumacher, C. Scotta, T. V. Shankey, A. Shemer, A. Simon, J. Spidlen, A. M. Stall, R. Stark, C. Stehle, M. Stein, T. Steinmetz, H. Stockinger, Y. Takahama, A. Tarnok, Z. Tian, G. Toldi, J. Tornack, E. Traggiai, J. Trotter, H. Ulrich, M. van der Braber, R. A. W. van Lier, M. Veldhoen, S. Vento -Asturias, P. Vieira, D. Voehringer, H. Volk, K. von Volkmann, A. Waisman, R. Walker, M. D. Ward, K. Warnatz, S. Warth, J. V. Watson, C. Watzl, L. Wegener, A. Wiedemann, J. Wienands, G. Willimsky, J. Wing, P. Wurst, L. Yu, A. Yue, Q. Zhang, Y. Zhao, S. Ziegler, and J. Zimmermann, Guidelines for the use of flow cytometry and cell sorting in immunological studies, Eur. J. Immunol., 2017, 47, 1584– 1797, doi:10.1002/eji.201646632.

[9] E. C. Jensen, Use of Fluorescent Probes: Their Effect on Cell Biology and Limitations, Anat. Rec., 2012, 295, 2031–2036, doi:10.1002/ar.22602.

[10] A. Ramoji, U. Neugebauer, T. Bocklitz, M. Foerster, M. Kiehntopf, M. Bauer, and J. Popp, Toward a Spectroscopic Hemogram: Raman Spectroscopic Differentiation of the Two Most Abundant Leukocytes from Peripheral Blood, Anal. Chem., 2012, 84, 5335–5342, doi:10.1021/ac3007363.

[11] B. Wollscheid, D. Bausch-Fluck, C. Henderson, R. O’Brien, M. Bibel, R. Schiess, R. Aebersold, and J. D. Watts, Mass-spectrometric identification and relative quantification of N-linked cell surface glycoproteins, Nat. Biotechnol., 2009, 27, 378–386, doi:10.1038/nbt.1532.

[12] Y. Park, C. Depeursinge, and G. Popescu, Quantitative phase imaging in biomedicine, Nat. Photonics, 2018, 12, 578–589, doi:10.1038/s41566-018-0253-x.

[13] R. Chandramohanadas, Y. Park, L. Lui, A. Li, D. Quinn, K. Liew, M. Diez-Silva, Y. Sung, M. Dao, C. T. Lim, P. R. Preiser, and S. Suresh, Biophysics of Malarial Parasite Exit from Infected Erythrocytes, PLoS One, 2011, 6, e20869, doi:10.1371/journal.pone.0020869.

[14] S. Koo, S. Jang, C. J. Park, Y.-U. Cho, and Y. Park, Unique Red Blood Cell Morphology Detected in a Patient with Myelodysplastic Syndrome by Three-dimensional Refractive Index Tomography, Lab. Med. Online, 2019, 9, 185–188.

[15] S. Oh, C. Lee, W. Yang, A. Li, A. Mukherjee, M. Basan, C. Ran, W. Yin, C. J. Tabin, and D. Fu, Protein and lipid mass concentration measurement in tissues by stimulated Raman scattering microscopy, Proceedings of the National Academy of Sciences, 2022, 119, e2117938119.

[16] S. Park, J. W. Ahn, Y. Jo, H.-Y. Kang, H. J. Kim, Y. Cheon, J. W. Kim, Y. Park, S. Lee, and K. Park, Label-Free Tomographic Imaging of Lipid Droplets in Foam Cells for Machine-Learning-Assisted Therapeutic Evaluation of Targeted Nanodrugs, ACS Nano, 2020, 14, 1856–1865, doi:10.1021/acsnano.9b07993.

[17] Y. Jo, S. Park, J. Jung, J. Yoon, H. Joo, M. Kim, S.-J. Kang, M. C. Choi, S. Y. Lee, and Y. Park, Holographic deep learning for rapid optical screening of anthrax spores, Sci. Adv., 2017, 3, e1700606, doi:10.1126/sciadv.1700606.

[18] F. Yi, I. Moon, and B. Javidi, Cell morphology-based classification of red blood cells using holographic imaging informatics, Biomed. Opt. Express, 2016, 7, 2385–2399, doi:10.1364/boe.7.002385.

[19] Q. T. K. Lai, K. C. M. Lee, K. K. Y. Wong, and K. K. Tsia, Label-free phytoplankton analysis by high-throughput quantitative phase imaging cytometry and machine learning, in 2019 Conference on Lasers and Electro-Optics Europe and European Quantum Electronics Conference, 2019, cl_5_1, https://opg.optica.org/abstract.cfm?uri=cleo_europe-2019-cl_5_1.

[20] D. Ryu, Y. Jo, J. Yoo, T. Chang, D. Ahn, Y. S. Kim, G. Kim, H.-S. Min, and Y. Park, Deep learning-based optical field screening for robust optical diffraction tomography, Sci. Rep., 2019, 9, 15239, doi:10.1038/s41598-019-51363-x.

[21] J. Yoon, Y. Jo, M. Kim, K. Kim, S. Lee, S.-J. Kang, and Y. Park, Identification of non-activated lymphocytes using three-dimensional refractive index tomography and machine learning, Sci. Rep., 2017, 7, 6654, doi:10.1038/s41598-017-06311-y.

[22] T. H. Nguyen, S. Sridharan, V. Macias, A. Kajdacsy-Balla, J. Melamed, M. N. Do, and G. Popescu, Automatic Gleason grading of prostate cancer using quantitative phase imaging and machine learning, J. Biomed. Opt., 2017, 22, 036015, doi:10.1117/1.jbo.22.3.036015.

[23] A. Cohen, M. Dudaie, I. Barnea, F. Borrelli, J. Běhal, L. Miccio, P. Memmolo, V. Bianco, P. Ferraro, and N. T. Shaked, Label-Free Imaging Flow Cytometry for Cell Classification Based on Multiple Interferometric Projections Using Deep Learning, Advanced Intelligent Systems, 2024, 6, 2300433, doi:10.1002/aisy.202300433.

[24] Y. Rivenson, Y. Zhang, H. Günaydın, D. Teng, and A. Ozcan, Phase recovery and holographic image reconstruction using deep learning in neural networks, Light Sci. Appl., 2018, 7, 17141, doi:10.1038/lsa.2017.141.

[25] A. Goy, K. Arthur, S. Li, and G. Barbastathis, Low Photon Count Phase Retrieval Using Deep Learning, Phys. Rev. Lett., 2018, 121, 243902, doi:10.1103/physrevlett.121.243902.

[26] Y. Rivenson, T. Liu, Z. Wei, Y. Zhang, K. de Haan, and A. Ozcan, PhaseStain: the digital staining of label-free quantitative phase microscopy images using deep learning, Light Sci. Appl., 2019, 8, 23, doi:10.1038/s41377-019-0129-y.

[27] G. Choi, D. Ryu, Y. Jo, Y. S. Kim, W. Park, H. Min, and Y. Park, Cycle-consistent deep learning approach to coherent noise reduction in optical diffraction tomography, Opt. Express, 2019, 27, 4927– 4943, doi:10.1364/oe.27.004927.

[28] T. Nguyen, Y. Xue, Y. Li, L. Tian, and G. Nehmetallah, Deep learning approach for Fourier ptychography microscopy, Opt. Express, 2018, 26, 26470–26484, doi:10.1364/oe.26.026470.

[29] Y. LeCun, Y. Bengio, and G. Hinton, Deep learning, Nature, 2015, 521, 436–444, doi:10.1038/nature14539.

[30] G. Kim, H. Hugonnet, K. Kim, J.-H. Lee, S. S. Lee, J. Ha, C. Lee, H. Park, K.-J. Yoon, Y. Shin, G. Csucs, I. Hitchcock, L. Mackinder, J. H. Kim, T. H. Hwang, S. Lee, P. O’Toole, B.-K. Koo, J. Guck, and Y. Park, Holotomography, Nat. Rev. Methods Primers, 2024, 4, 51, doi:10.1038/s43586-024-00327-1.

[31] S. H. Karandikar, C. Zhang, A. Meiyappan, I. Barman, C. Finck, P. K. Srivastava, and R. Pandey, Reagent-Free and Rapid Assessment of T Cell Activation State Using Diffraction Phase Microscopy and Deep Learning, Anal. Chem., 2019, 91, 3405–3411, doi:10.1021/acs.analchem.8b04895.

[32] L. Zhao, L. Tang, M. S. Greene, Y. Sa, W. Wang, J. Jin, H. Hong, J. Q. Lu, and X.-H. Hu, Deep Learning of Morphologic Correlations To Accurately Classify CD4+ and CD8+ T Cells by Diffraction Imaging Flow Cytometry, Anal. Chem., 2022, 94, 1567–1574, doi:10.1021/acs.analchem.1c03337.

[33] V. Gorti, C. E. Serafini, A. D. Silva Trenkle, K. McCubbins, I. LeCompte, G. A. Kwong, and F. E. Robles, Nondestructive, High-Resolution T Cell Characterization and Subtyping via Deep-UV Microscopy, BME Front., 2026, 7, 0227, doi:10.34133/bmef.0227.

[34] D. Yagoda-Aharoni, E. Dotan, M. Dudaie, and N. T. Shaked, Label-Free Holographic Imaging Flow Cytometry With Deep-Learning-Based Detection and Classification of Thousands of Cells Per Second, Cytometry A, 2026, 109, 89–97, doi:10.1002/cytoa.70008.

[35] S. Sun, J. Pang, J. Shi, S. Yi, and W. Ouyang, Fishnet: A versatile backbone for image, region, and pixel level prediction, Advances in neural information processing systems, 2018, 31, https://proceedings.neurips.cc/paper/2018/hash/75fc093c0ee742f6dddaa13fff98f104-Abstract.html.

[36] G. Huang, Z. Liu, L. Van Der Maaten, and K. Q. Weinberger, Densely connected convolutional networks, in Proceedings of the IEEE Conference on Computer Vision and Pattern Recognition (CVPR), 2017, 4700–4708, doi:10.1109/CVPR.2017.243.

[37] J. Hu, L. Shen, S. Albanie, G. Sun, and E. Wu, Squeeze-and-Excitation Networks, IEEE Trans. Pattern Anal. Mach. Intell., 2020, 42, 2011–2023, doi:10.1109/tpami.2019.2913372.

[38] S. Shin, K. Kim, J. Yoon, and Y. Park, Active illumination using a digital micromirror device for quantitative phase imaging, Opt. Lett., 2015, 40, 5407–5410, doi:10.1364/ol.40.005407.

[39] K. Lee, K. Kim, G. Kim, S. Shin, and Y. Park, Time-multiplexed structured illumination using a DMD for optical diffraction tomography, Opt. Lett., 2017, 42, 999–1002, doi:10.1364/ol.42.000999.

[40] M. Takeda, H. Ina, and S. Kobayashi, Fourier-transform method of fringe-pattern analysis for computer-based topography and interferometry, J. Opt. Soc. Am., 1982, 72, 156–160, doi:10.1364/JOSA.72.000156.

[41] S. K. Debnath, and Y. Park, Real-time quantitative phase imaging with a spatial phase-shifting algorithm, Opt. Lett., 2011, 36, 4677–4679, doi:10.1364/ol.36.004677.

[42] E. Wolf, Three-dimensional structure determination of semi-transparent objects from holographic data, Opt. Commun., 1969, 1, 153–156, doi:10.1016/0030-4018(69)90052-2.

[43] J. Lim, K. Lee, K. H. Jin, S. Shin, S. Lee, Y. Park, and J. C. Ye, Comparative study of iterative reconstruction algorithms for missing cone problems in optical diffraction tomography, Opt. Express, 2015, 23, 16933–16948, doi:10.1364/oe.23.016933.

[44] V. Lauer, New approach to optical diffraction tomography yielding a vector equation of diffraction tomography and a novel tomographic microscope, J. Microsc., 2002, 205, 165–176, doi:10.1046/j.0022-2720.2001.00980.x.

[45] C. Park, S. Shin, and Y. Park, Generalized quantification of three-dimensional resolution in optical diffraction tomography using the projection of maximal spatial bandwidths, J. Opt. Soc. Am. A, 2018, 35, 1891–1898, doi:10.1364/josaa.35.001891.

[46] G. Popescu, Y. Park, N. Lue, C. Best-Popescu, L. Deflores, R. R. Dasari, M. S. Feld, and K. Badizadegan, Optical imaging of cell mass and growth dynamics, Am. J. Physiol. Cell Physiol., 2008, 295, C538–C544, doi:10.1152/ajpcell.00121.2008.

[47] R. Barer, Determination of Dry Mass, Thickness, Solid and Water Concentration in Living Cells, Nature, 1953, 172, 1097–1098, doi:10.1038/1721097a0.

[48] D. Ryu, J. Kim, D. Lim, H.-S. Min, I. Y. Yoo, D. Cho, and Y. Park, Label-Free White Blood Cell Classification Using Refractive Index Tomography and Deep Learning, BME Front., 2021, 2021, 9893804, doi:10.34133/2021/9893804.

[49] M. Lee, Y.-H. Lee, J. Song, G. Kim, Y. Jo, H. Min, C. H. Kim, and Y. Park, Deep-learning-based three-dimensional label-free tracking and analysis of immunological synapses of CAR-T cells, Elife, 2020, 9, e49023, doi:10.7554/elife.49023.

[50] M. Sung, J. H. Kim, H.-S. Min, S. Jang, J. Hong, B. K. Choi, J. Shin, K. S. Chung, and Y. R. Park, Three-dimensional label-free morphology of CD8+ T cells as a sepsis biomarker, Light: Science & Applications, 2023, 12, 265, doi:10.1038/s41377-023-01309-w.

[51] R. R. Selvaraju, M. Cogswell, A. Das, R. Vedantam, D. Parikh, and D. Batra, Grad-cam: Visual explanations from deep networks via gradient-based localization, 2017, 618–626.

[52] G. Montavon, W. Samek, and K.-R. Müller, Methods for interpreting and understanding deep neural networks, Digit. Signal Process., 2018, 73, 1–15, doi:10.1016/j.dsp.2017.10.011.

[53] T. Ching, D. S. Himmelstein, B. K. Beaulieu-Jones, A. A. Kalinin, B. T. Do, G. P. Way, E. Ferrero, P.-M. Agapow, M. Zietz, M. M. Hoffman, W. Xie, G. L. Rosen, B. J. Lengerich, J. Israeli, J. Lanchantin, S. Woloszynek, A. E. Carpenter, A. Shrikumar, J. Xu, E. M. Cofer, C. A. Lavender, S. C. Turaga, A. M. Alexandari, Z. Lu, D. J. Harris, D. DeCaprio, Y. Qi, A. Kundaje, Y. Peng, L. K. Wiley, M. H. S. Segler, S. M. Boca, S. J. Swamidass, A. Huang, A. Gitter, and C. S. Greene, Opportunities and obstacles for deep learning in biology and medicine, J. R. Soc. Interface, 2018, 15, 20170387, doi:10.1098/rsif.2017.0387.

