## Supplementary Information for "Three-Dimensional Refractive Index Patterns Distinguish CD57-Defined Subsets of Human Memory T Cells"

#### Supplementary Methods

##### S1 Reagents and reference populations

Peripheral blood mononuclear cells were isolated by Ficoll density-gradient centrifugation (Corning, Inc., USA). T cells were enriched by magnetic negative selection using the Pan T Cell Isolation Kit (Miltenyi Biotec, USA). The antibody panel comprised BV421 anti-CD4 (RPA-T4, BD Biosciences, USA), APC-H7 anti-CD8 (SK1, BD Biosciences), PE-Cy7 anti-CD3 (UCHT1, eBioscience, USA), APC anti-CD45RA (HI100, eBioscience), PE anti-CD57 (HNK-1, BioLegend, USA), and FITC anti-CCR7 (150503, R&D Systems, USA). Dead cells were excluded using 7-aminoactinomycin D (7-AAD; BD Biosciences).

Fluorescence-activated cell sorting (FACS) on a FACSAria II instrument (BD Biosciences) yielded CD4+ T cells, naïve (CCR7+CD45RA+) CD8+ T cells, and CD57− and CD57+ memory CD8+ T cells. The memory fraction was defined as CCR7−CD45RA+/−. These sorted populations supplied the reference labels for classification; the convolutional neural network (CNN) received refractive index (RI) volumes without fluorescence inputs.

##### S2 Optical configuration and reconstruction

The HT-2H microscope (Tomocube, Inc., Republic of Korea) implements coherent holotomography (HT) using a laser-based Mach–Zehnder interferometer (Supplementary Fig. 1). A 2 × 2 fiber coupler divided the output of a coherent 532 nm laser (vacuum wavelength; Samba, Cobolt Inc., Sweden) into sample and reference beams. The sample beam passed through a 4-f relay (lenses L1–L3) and a digital micromirror device (DMD; DLPLCR6500EVM, Texas Instruments Inc., USA), which displayed digital grating patterns to control the illumination angle.

Lens L4 and a water-immersion condenser objective (UPLSAPO 60XW, Olympus Inc.; numerical aperture 1.2) projected the first diffraction order onto the sample at 49 angles. A matched collection objective captured the scattered light, which interfered with the reference beam at the camera (FL3-U3-13Y3M-C, Point Gray Research Inc.). A linear polarizer increased fringe contrast. The full 49-hologram sequence required approximately 0.1 s per cell.

Complex fields recovered from the holograms were combined by Fourier-diffraction-theorem inversion in TomoStudio. Forty iterations of non-negativity regularization suppressed missing-cone artifacts. The lateral and axial resolutions were 110 nm and 355 nm, respectively. The reconstruction methods and their references are described in the main Methods.

##### S3 Feature extraction and classifier training

The k-nearest neighbor (k-NN) baseline used cellular volume (CV), surface area (SA), sphericity, protein density (PD), and dry mass (DM) evaluated at 20 RI thresholds from 1.340 to 1.378 in increments of 0.002. This yielded 100 features per cell (Supplementary Fig. 2). Several values of k were screened; the reported classifier used k = 5 and Euclidean distance. Binary support vector machines (SVMs) were implemented using MATLAB fitcsvm and multiclass SVMs using fitcecoc. Kernel type, box constraint, and kernel scale were tuned for each task by cross-validation on the training partition.

The three-dimensional (3-D) CNN used a FishNet-inspired tail–body–head architecture with dense connectivity and squeeze-and-excitation channel attention. Supplementary Table 1 retains the detailed layer specification. Batch-normalization momentum was 0.1, and the LeakyReLU negative slope was 0.01. The decision block projected 2,112 channels to 1,056 and then to four outputs. The structural connections and output sequence are shown in Fig. 3 of the main manuscript.

Training used the adaptive moment estimation (Adam) optimizer with a learning rate of 0.001, a reported momentum parameter of 0.9, and a learning-rate decay of 0.9999. The batch size was 8. Data augmentation comprised random translation, random cropping, elastic transformation, and additive Gaussian noise. The implementation used PyTorch 1.0.1. Training ran for 500 epochs and required approximately 20 h on a workstation with four NVIDIA GeForce GTX 1080 Ti graphics processing units and an Intel Xeon E5-2600 central processing unit. Validation accuracy selected the checkpoint for testing.

The four-class CNN cohort comprised 1,354 cells: 482 CD4+, 334 naïve CD8+, 252 CD57− memory CD8+, and 286 CD57+ memory CD8+. The reported 6:2:2 cell-level partition included 814 training and 270 held-out-test cells; subtraction from the total gives 270 validation cells. The binary models had separate task-specific partitions: 698 training and 87 test cells for naïve versus memory CD8+, and 430 training and 54 test cells for CD57− versus CD57+ memory CD8+. The original analyst recalled an approximately 8:1:1 split for the binary tasks, but exact validation counts and donor membership were not recoverable. Test counts and performance are summarized in Supplementary Tables 3 and 4.

The k-NN and SVM summaries each used 210, 108, and 107 test cells for the four-class, naïve-versus-memory, and CD57 tasks, respectively. The corresponding CNN tests used 270, 87, and 54 cells. These model-specific partitions differ, so cross-model accuracies are descriptive comparisons. The DenseNet result is available only as a four-class accuracy range of 78–83%; the associated settings and partition membership were not specified in the recovered summary (Supplementary Table 2).

#### S4 Morphometry cohorts and visualization

The morphometry cohort in main Table 1 comprised 246 CD4+, 270 naïve CD8+, 252 CD57− memory CD8+, and 286 CD57+ memory CD8+ cells. It differs from the 1,354-cell CNN learning cohort. Cell-level correspondence between these cohorts could not be established from the available records.

The individual observations in main Fig. 2b were reconstructed from marker coordinates in the original vector plots. CV, sphericity, PD, and DM each contain 246 plotted points per class. SA contains 245 CD4+, 245 naïve CD8+, 246 CD57− memory CD8+, and 246 CD57+ memory CD8+ points. These reconstructed values are approximate and represent the available plotted subsets rather than the complete Table 1 cohort. Boxplots show the median, interquartile range, and observations within 1.5 times that range; remaining points are displayed as outliers. The original analysis used MATLAB built-in statistical functions, but the underlying cell-level records were unavailable for reproducing inferential comparisons. The current plots and discussion therefore describe the observed distributions without significance annotations.

For main Fig. 4d, 120 tomograms were randomly selected, 30 from each FACS-defined class. Their four-component final CNN output vectors were embedded in two dimensions with MATLAB R2018b's t-distributed stochastic neighbor embedding (t-SNE) implementation. The embedding provides a qualitative representation of output similarity; the symbols denote reference identities. The separate feature-vector embeddings for k-NN are retained in Supplementary Fig. 4.

##### S5 Mixed-sample fractions and model-development history

The mixed-sample comparison in main Fig. 5 covers three donors measured on three experimental days. Archived slides document 1,100 unsorted Day 1 cells assigned assumed CD4+ labels because the corresponding FACS sample was enriched for CD4+ cells. The slides allocate 500 cells to training, 350 to validation, and 250 to testing. The original analyst recalled including these additional data when training the mixed-sample model. Day 1 therefore contributed to model development; the overlap between the development cells and those used for the plotted fractions could not be verified.

Supplementary Table 5 reproduces the fractions used in Fig. 5. The selected summary dated April 25, 2026 gives identical FACS values for Days 2 and 3. Values are retained as reported, without renormalization. The analyzed cell counts, fraction denominators, and inclusion criteria for the two methods were unavailable, so the entries support a descriptive comparison of reported fractions. They cannot be used to calculate independent composition accuracy or to assign the discrepancies to a specific source of error.

#### Supplementary Figure 1

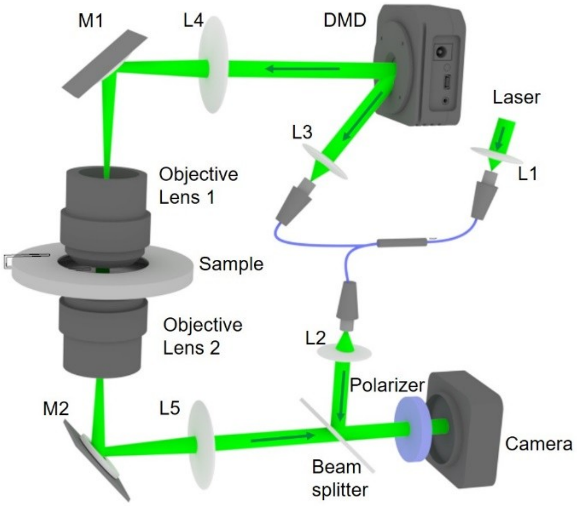


Supplementary Figure 1. Optical configuration of the HT-2H coherent holotomography (HT) system. A coherent 532 nm laser supplies the sample and reference arms of a Mach–Zehnder interferometer. A digital micromirror device (DMD) controls the sample illumination angle. L1–L5, lenses; M1 and M2, mirrors. The beam path and components are described in Supplementary Methods S2.

#### Supplementary Figure 2

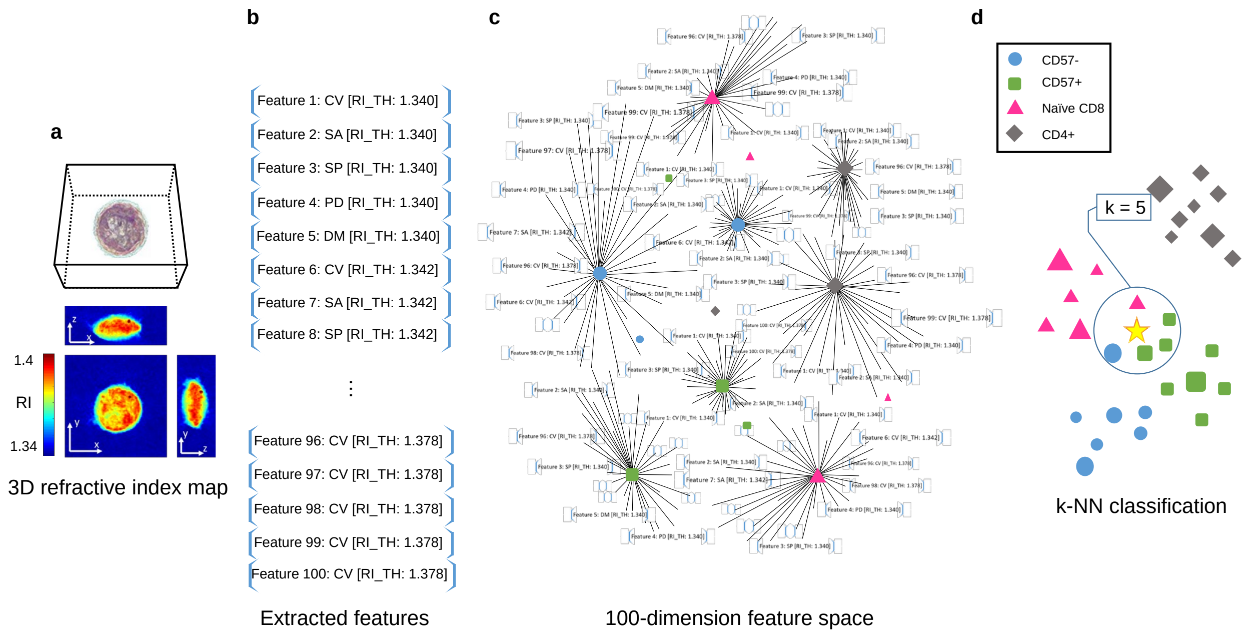


Supplementary Figure 2. Feature extraction and k-nearest neighbor (k-NN) classification. a, Input three-dimensional (3-D) refractive index (RI) tomogram. b, Cellular volume (CV), surface area (SA), sphericity (SP), protein density (PD), and dry mass (DM) are calculated at 20 RI thresholds (RI-TH), producing a 100-component vector. c, The vectors define a Euclidean feature space. d, The test cell (yellow star) is assigned the majority label among its five nearest neighbors. In this schematic example, three of the five neighbors are CD57+ memory CD8+ cells (green squares), so the test cell receives that label. Blue circles denote CD57− memory CD8+ cells, pink triangles naïve CD8+ cells, and gray diamonds CD4+ cells.

### Supplementary Figure 3

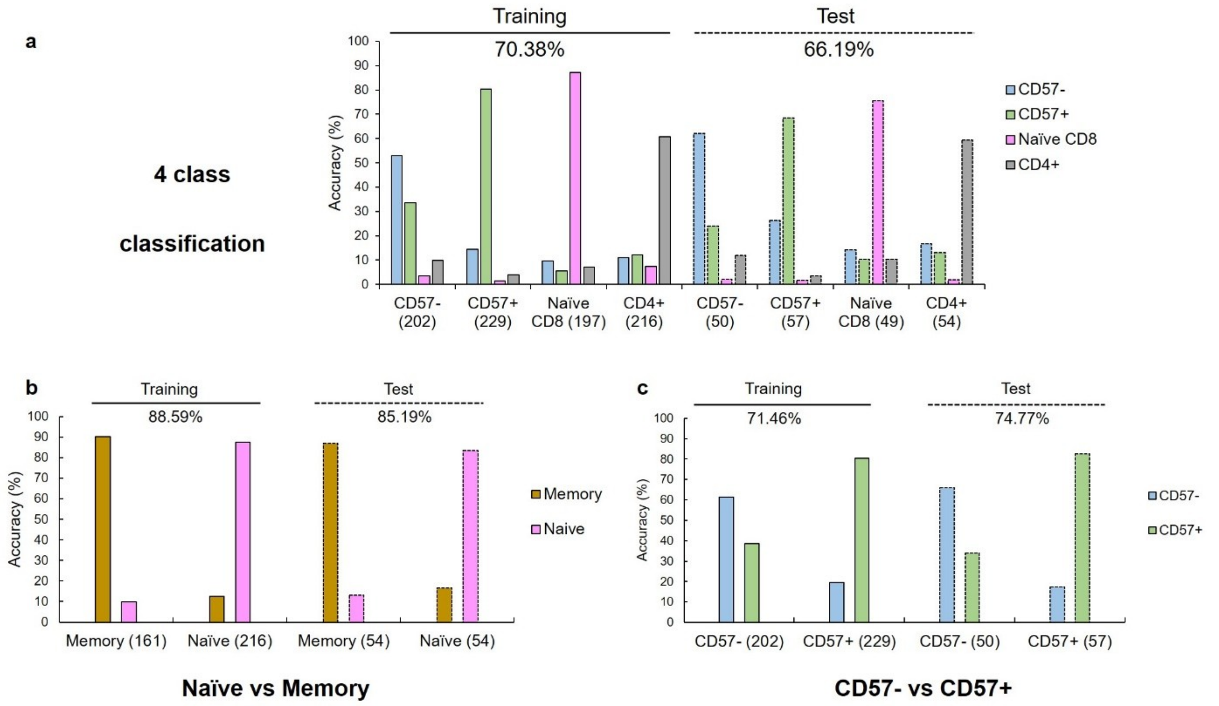


Supplementary Figure 3. Training and test results for k-nearest neighbor (k-NN) classification. a, Four T-cell classes. b, Naïve versus memory CD8+ cells. c, CD57− versus CD57+ memory CD8+ cells. Each group on the horizontal axis denotes a reference population, with its cell count in parentheses; colored bars show the distribution of predicted classes. Test accuracies are 66.19% (139/210), 85.19% (92/108), and 74.77% (80/107) for panels a–c, respectively. CD57− and CD57+ denote the memory CD8+ subsets.

#### Supplementary Figure 4

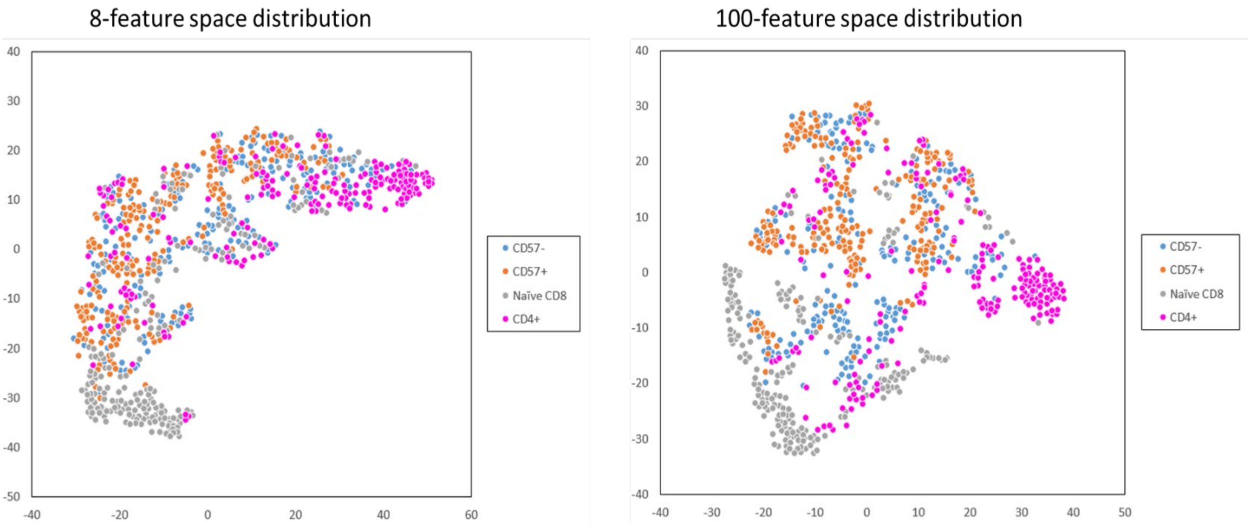


Supplementary Figure 4. Two-dimensional t-distributed stochastic neighbor embedding (t-SNE) of the feature vectors used in the k-nearest neighbor (k-NN) analyses. The archived plots show 8-component vectors (left) and 100-component vectors (right). Colors identify CD57− memory CD8+ (blue), CD57+ memory CD8+ (orange), naïve CD8+ (gray), and CD4+ (magenta) cells. Axes are arbitrary embedding coordinates.

#### Supplementary Figure 5

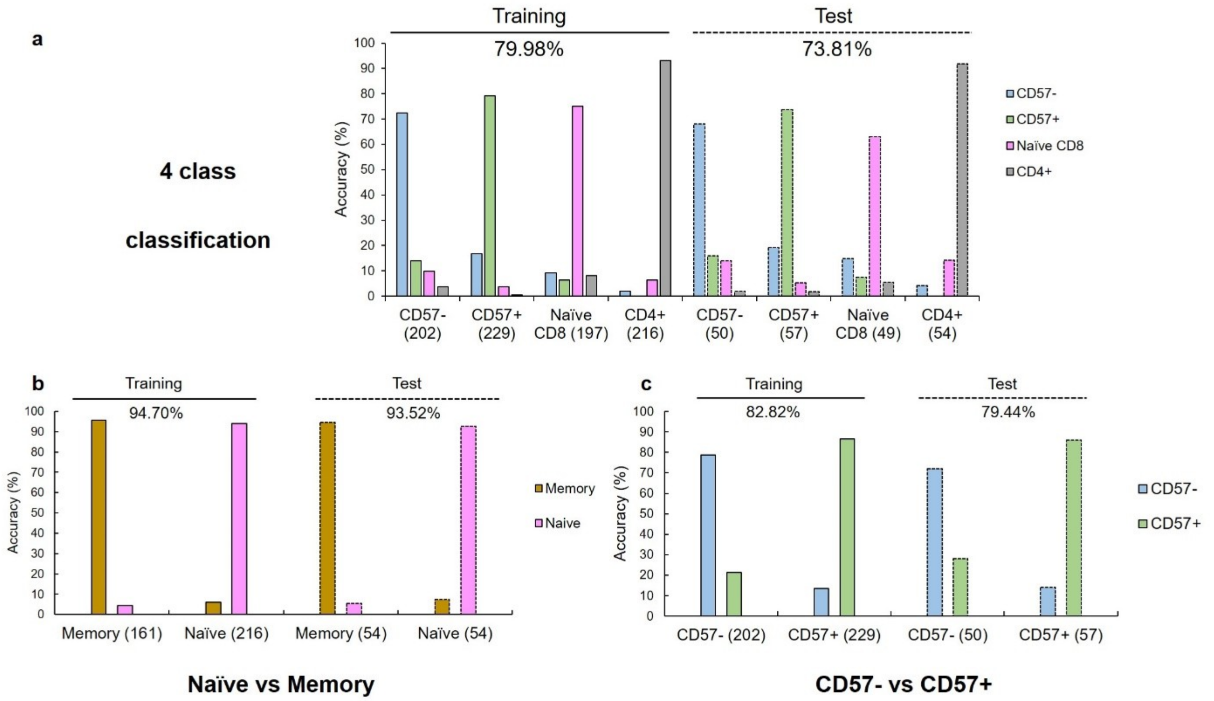


Supplementary Figure 5. Training and test results for support vector machine (SVM) classification. a, Four T-cell classes. b, Naïve versus memory CD8+ cells. c, CD57− versus CD57+ memory CD8+ cells. Reference populations and cell counts appear on the horizontal axis; colored bars show predicted-class distributions. Test accuracies are 73.81% (155/210), 93.52% (101/108), and 79.44% (85/107) for panels a–c, respectively. Model-specific test partitions differ from those of the convolutional neural networks (Supplementary Table 3).

### Supplementary Table 1

Supplementary Table 1. Layer specification of the 3-D CNN. The two-page specification lists the convolutional stem, tail, bridge, body, head, and classifier, with channel counts and layer parameters. Conv3d, three-dimensional convolution; BatchNorm3d, three-dimensional batch normalization; MaxPool3d, three-dimensional max pooling; LeakyReLU, leaky rectified linear unit; NN, nearest-neighbor upsampling; ch, channels.

| class 1 | class 2 | class3 | layer | function | input channel | out channel | kernel | stride | padding | momentum | negative slope | dilation | ch 1 | ch2 | ch3 | upsample factor | mode |
|---|---|---|---|---|---|---|---|---|---|---|---|---|---|---|---|---|---|
| Stem | | sequential | | Conv3d | 1 | 32 | 3 | 2 | 1 | | | | | | | | |
| | | | | BatchNorm3d | | 32 | | | | 0.1 | | | | | | | |
| | | | | LeakyReLU | 32 | 32 | | | | | 0.01 | | | | | | |
| | | sequential | | Conv3d | 32 | 32 | 3 | 1 | 1 | | | | | | | | |
| | | | | BatchNorm3d | | | | | | 0.1 | | | | | | | |
| | | | | LeakyReLU | | | | | | | 0.01 | | | | | | |
| | | sequential | | Conv3d | 32 | 64 | 3 | 1 | 1 | | | | | | | | |
| | | | | BatchNorm3d | | | | | | 0.1 | | | | | | | |
| | | | | LeakyReLU | | | | | | | 0.01 | | | | | | |
| | | pool | | MaxPool3d | | | 3 | 2 | 1 | | | 1 | | | | | |
| Tail | Fishtail1 | ResBlock 1 | | Residual Block | | | | | | | | | 64 | 32 | 128 | | |
| | | | Shortcut | BatchNorm3d | | | | | | 0.1 | | | | | | | |
| | | | | LeakyReLU | | | | | | | 0.01 | | | | | | |
| | | | | Conv3d | 64 | 128 | 1 | 1 | | | | | | | | | |
| | | ResBlock 2 | | Residual Block | | | | | | | | | 128 | 32 | 128 | | |
| | | Pooling | | MaxPool3d | | | 2 | 2 | 0 | | | 1 | | | | | |
| | FishTail2 | ResBlock 1 | | Residual Block | | | | | | | | | 128 | 64 | 256 | | |
| | | | Shortcut | BatchNorm3d | | | | | | 0.1 | | | | | | | |
| | | | | LeakyReLU | | | | | | | 0.01 | | | | | | |
| | | | | Conv3d | 128 | 256 | 1 | 1 | | | | | | | | | |
| | | ResBlock 2 | | Residual Block | | | | | | | | | 256 | 64 | 256 | | |
| | | ResBlock 3 | | Residual Block | | | | | | | | | 256 | 64 | 256 | | |
| | | ResBlock 4 | | Residual Block | | | | | | | | | 256 | 64 | 256 | | |
| | | ResBlock 5 | | Residual Block | | | | | | | | | 256 | 64 | 256 | | |
| | | ResBlock 6 | | Residual Block | | | | | | | | | 256 | 64 | 256 | | |
| | | ResBlock 7 | | Residual Block | | | | | | | | | 256 | 64 | 256 | | |
| | | ResBlock 8 | | Residual Block | | | | | | | | | 256 | 64 | 256 | | |
| | | Pooling | | MaxPool3d | | | 2 | 2 | 0 | | | 1 | | | | | |
| | FishTail3 | ResBlock 1 | | Residual Block | | | | | | | | | 256 | 128 | 512 | | |
| | | | Shortcut | BatchNorm3d | | | | | | 0.1 | | | | | | | |
| | | | | LeakyReLU | | | | | | | 0.01 | | | | | | |
| | | | | Conv3d | 256 | 512 | 1 | 1 | | | | | | | | | |
| | | ResBlock 3 | | Residual Block | | | | | | | | | 512 | 158 | 512 | | |
| | | ResBlock 4 | | Residual Block | | | | | | | | | 512 | 158 | 512 | | |
| | | ResBlock 5 | | Residual Block | | | | | | | | | 512 | 158 | 512 | | |
| | | ResBlock 6 | | Residual Block | | | | | | | | | 512 | 158 | 512 | | |
| | | ResBlock 7 | | Residual Block | | | | | | | | | 512 | 158 | 512 | | |
| | | ResBlock 8 | | Residual Block | | | | | | | | | 512 | 158 | 512 | | |
| | | ResBlock 9 | | Residual Block | | | | | | | | | 512 | 158 | 512 | | |
| | | ResBlock 10 | | Residual Block | | | | | | | | | 512 | 158 | 512 | | |
| | | ResBlock 11 | | Residual Block | | | | | | | | | 512 | 158 | 512 | | |
| | | ResBlock 12 | | Residual Block | | | | | | | | | 512 | 158 | 512 | | |
| | | ResBlock 13 | | Residual Block | | | | | | | | | 512 | 158 | 512 | | |
| | | ResBlock 14 | | Residual Block | | | | | | | | | 512 | 158 | 512 | | |
| | | ResBlock 15 | | Residual Block | | | | | | | | | 512 | 158 | 512 | | |
| | | ResBlock 16 | | Residual Block | | | | | | | | | 512 | 158 | 512 | | |
| | | Pooling | | MaxPool3d | | | 2 | 2 | 0 | | | 1 | | | | | |
| Bridge | stem | Sequential | | BatchNorm3d | | | | | | 0.1 | | | | | | | |
| | | | | LeakyReLU | | | | | | | 0.01 | | | | | | |
| | | | | Conv3d | 512 | 256 | 1 | 1 | | | | | | | | | |
| | | | | BatchNorm3d | | | | | | 0.1 | | | | | | | |
| | | | | LeakyReLU | | | | | | | 0.01 | | | | | | |
| | | | | Conv3d | 256 | 1024 | 1 | 1 | | | | | | | | | |
| | Sequential | | ResBlock 1 | Residual Block | | | | | | | | | 1024 | 128 | 512 | | |
| | | | Shortcut | BatchNorm3d | | | | | | 0.1 | | | | | | | |
| | | | | LeakyReLU | | | | | | | 0.01 | | | | | | |
| | | | | Conv3d | 1025 | 512 | 1 | 1 | | | | | | | | | |
| | | | ResBlock 2 | Residual Block | | | | | | | | | 512 | 128 | 512 | | |
| | | | ResBlock 3 | Residual Block | | | | | | | | | 512 | 128 | 512 | | |
| | | | ResBlock 4 | Residual Block | | | | | | | | | 512 | 128 | 512 | | |
| | | | ResBlock 5 | Residual Block | | | | | | | | | 512 | 128 | 512 | | |
| | | | ResBlock 6 | Residual Block | | | | | | | | | 512 | 128 | 512 | | |
| | | | ResBlock 7 | Residual Block | | | | | | | | | 512 | 128 | 512 | | |
| | | | ResBlock 8 | Residual Block | | | | | | | | | 512 | 128 | 512 | | |

### Supplementary Table 1 (continued)

| | | | | | | | | | | | | | | | | | |
|---|---|---|---|---|---|---|---|---|---|---|---|---|---|---|---|---|---|
| | se block | | | BatchNorm3d | | | | | | 0.1 | | | | | | | |
| | | | | LeakyReLU | | | | | | | 0.01 | | | | | | |
| | | | | AdaptiveAverage Pooling3d | | 1 | | | | | | | | | | | |
| | | | | Conv3d | 1024 | 32 | 1 | 1 | | | | | | | | | |
| | | | | LeakyReLU | | | | | | | 0.01 | | | | | | |
| | | | | Conv3d | 32 | 512 | 1 | 1 | | | | | | | | | |
| | | | | Sigmoid | | | | | | | | | | | | | |
| Body | FishBody1 | ResBlock 1 | | Residual Block | | | | | | | | | 512 | 128 | 512 | | |
| | | ResBlock 2 | | Residual Block | | | | | | | | | 512 | 128 | 512 | | |
| | | ResBlock 3 | | Residual Block | | | | | | | | | 512 | 128 | 512 | | |
| | | ResBlock 4 | | Residual Block | | | | | | | | | 512 | 128 | 512 | | |
| | | upsample | | Upsampling | | | | 0 | | | | | | | | 2 | NN |
| | | TransferBlock | ResBlock 1 | Residual Block | | | | | | | | | 256 | 64 | 256 | | |
| | | | ResBlock 2 | Residual Block | | | | | | | | | 256 | 64 | 256 | | |
| | FishBody2 | ResBlock 1 | | Residual Block | | | | | | | | | 768 | 96 | 384 | | |
| | | ResBlock 2 | | Residual Block | | | | | | | | | 384 | 96 | 384 | | |
| | | ResBlock 3 | | Residual Block | | | | | | | | | 384 | 96 | 384 | | |
| | | ResBlock4 | | Residual Block | | | | | | | | | 384 | 96 | 384 | | |
| | | upsample | | Upsampling | | | | | | | | | | | | 2 | NN |
| | | TransferBlock | ResBlock 1 | Residual Block | | | | | | | | | 128 | 32 | 128 | | |
| | | | ResBlock 2 | Residual Block | | | | | | | | | 128 | 32 | 128 | | |
| | FishBody3 | ResBlock 1 | | Residual Block | | | | | | | | | 512 | 64 | 256 | | |
| | | ResBlock 2 | | Residual Block | | | | | | | | | 256 | 64 | 256 | | |
| | | ResBlock 3 | | Residual Block | | | | | | | | | 256 | 64 | 256 | | |
| | | ResBlock4 | | Residual Block | | | | | | | | | 256 | 64 | 256 | | |
| | | upsample | | Upsampling | | | | 0 | | | | | | | | 2 | NN |
| | | TransferBlock | ResBlock 1 | Residual Block | | | | | | | | | 64 | 16 | 64 | | |
| | | | ResBlock 2 | Residual Block | | | | | | | | | 64 | 16 | 64 | | |
| | | | ResBlock 3 | Residual Block | | | | | | | | | 64 | 16 | 64 | | |
| | | | ResBlock 4 | Residual Block | | | | | | | | | 64 | 16 | 64 | | |
| Head | FishHead1 | ResBlock 1 | | Residual Block | | | | | | | | | 320 | 80 | 320 | | |
| | | ResBlock 2 | | Residual Block | | | | | | | | | 320 | 80 | 320 | | |
| | | ResBlock 3 | | Residual Block | | | | | | | | | 320 | 80 | 320 | | |
| | | ResBlock 4 | | Residual Block | | | | | | | | | 320 | 80 | 320 | | |
| | | Pooling | | MaxPool3d | | | 2 | 2 | 0 | | | 1 | | | | | |
| | | TransferBlock | ResBlock 1 | Residual Block | | | | | | | | | 512 | 128 | 512 | | |
| | | | ResBlock 2 | Residual Block | | | | | | | | | 512 | 128 | 512 | | |
| | FishHead2 | ResBlock 1 | | Residual Block | | | | | | | | | 832 | 208 | 832 | | |
| | | ResBlock 2 | | Residual Block | | | | | | | | | 832 | 208 | 832 | | |
| | | ResBlock 3 | | Residual Block | | | | | | | | | 832 | 208 | 832 | | |
| | | ResBlock 4 | | Residual Block | | | | | | | | | 832 | 208 | 832 | | |
| | | Pooling | | MaxPool3d | | | 2 | 2 | 0 | | | 1 | | | | | |
| | | TransferBlock | ResBlock 1 | Residual Block | | | | | | | | | 768 | 192 | 768 | | |
| | | | ResBlock 2 | Residual Block | | | | | | | | | 768 | 192 | 768 | | |
| | FishHead3 | ResBlock 1 | | Residual Block | | | | | | | | | 1600 | 400 | 1600 | | |
| | | ResBlock 2 | | Residual Block | | | | | | | | | 1600 | 400 | 1600 | | |
| | | ResBlock 3 | | Residual Block | | | | | | | | | 1600 | 400 | 1600 | | |
| | | ResBlock 4 | | Residual Block | | | | | | | | | 1600 | 400 | 1600 | | |
| | | ResBlock 5 | | Residual Block | | | | | | | | | 1600 | 400 | 1600 | | |
| | | ResBlock 6 | | Residual Block | | | | | | | | | 1600 | 400 | 1600 | | |
| | | ResBlock 7 | | Residual Block | | | | | | | | | 1600 | 400 | 1600 | | |
| | | ResBlock 8 | | Residual Block | | | | | | | | | 1600 | 400 | 1600 | | |
| | | Pooling | | MaxPool3d | | | 2 | 2 | 0 | | | 1 | | | | | |
| | | TransferBlock | ResBlock 1 | Residual Block | | | | | | | | | 512 | 128 | 512 | | |
| | | | ResBlock 2 | Residual Block | | | | | | | | | 512 | 128 | 512 | | |
| | | | ResBlock 3 | Residual Block | | | | | | | | | 512 | 128 | 512 | | |
| | | | ResBlock 4 | Residual Block | | | | | | | | | 512 | 128 | 512 | | |
| Classifier | | | | BatchNorm3d | | 2112 | | | | 0.1 | | | | | | | |
| | | | | LeakyReLU | | | | | | | 0.01 | | | | | | |
| | | | | Conv3d | 2112 | 1056 | 1 | 1 | 1 | | | | | | | | |
| | | | | BatchNorm3d | | 1056 | | | | 0.1 | | | | | | | |
| | | | | LeakyReLU | | | | | | | 0.01 | | | | | | |
| | | | | AdaptiveAvgPool 3d | | 1 | 1 | 1 | 1 | | | | | | | | |
| | | | | Conv3d | 1056 | 4 | | | | | | | | | | | |

**Supplementary Table 2**

| | K-NN (ML) | SVM (ML) | DenseNet (DL) | FishNet (DL) |
|---|---|---|---|---|
| 4 classes | 66.19% | 73.81% | 78 - 83 % | 97.04% |
| Naïve / Memory | 85.19% | 93.52% | | 97.70% |
| CD57- / CD57+ | 74.77% | 79.44% | | 94.44% |

Supplementary Table 2. Reported test accuracies by classifier. k-NN, k-nearest neighbors; SVM, support vector machine; ML, machine learning; DL, deep learning. FishNet denotes the FishNet-inspired three-dimensional convolutional neural network used here. The CD57 task concerns the two memory CD8+ subsets. DenseNet is reported only for the four-class task; the recovered summary does not identify the settings or partitions associated with its 78–83% range. The test partitions differ among models. Exact counts for the k-NN, SVM, and CNN results appear in Supplementary Table 3.

#### Supplementary Table 3. Test results and task-specific partitions

| Task | Classifier | Correct | Test cells | Accuracy (%) |
|---|---|---|---|---|
| Four classes | k-NN | 139 | 210 | 66.19 |
| Four classes | SVM | 155 | 210 | 73.81 |
| Four classes | CNN | 262 | 270 | 97.04 |
| Naïve vs memory CD8+ | k-NN | 92 | 108 | 85.19 |
| Naïve vs memory CD8+ | SVM | 101 | 108 | 93.52 |
| Naïve vs memory CD8+ | CNN | 85 | 87 | 97.70 |
| Memory CD8+ CD57− vs CD57+ | k-NN | 80 | 107 | 74.77 |
| Memory CD8+ CD57− vs CD57+ | SVM | 85 | 107 | 79.44 |
| Memory CD8+ CD57− vs CD57+ | CNN | 51 | 54 | 94.44 |

k-NN, k-nearest neighbors; SVM, support vector machine; CNN, convolutional neural network. Counts are correctly classified cells and total test cells. Model-specific test partitions differ.

| CNN task | Training cells | Validation cells | Test cells |
|---|---|---|---|
| Four classes | 814 | 270* | 270 |
| Naïve vs memory CD8+ | 698 | Not recovered | 87 |
| Memory CD8+ CD57− vs CD57+ | 430 | Not recovered | 54 |

*Four-class validation count inferred as 1,354 − 814 − 270 = 270. The original analyst recalled an approximately 8:1:1 split for binary models; exact validation counts and donor membership were unavailable. These are separate cell-level evaluations.

#### Supplementary Table 4. CNN training accuracy and four-class test recall

| Task | Correct training cells | Training cells | Accuracy (%) |
| --- | --- | --- | --- |
| Four classes | 802 | 814 | 98.53 |
| Naïve vs memory CD8+ | 697 | 698 | 99.86 |
| Memory CD8+ CD57− vs CD57+ | 418 | 430 | 97.21 |

Training accuracies refer to the reported models. Held-out-test accuracies are listed in Supplementary Table 3.

| Reference class | Correct test cells | Test cells | Recall (%) |
| --- | --- | --- | --- |
| CD4+ | 94 | 96 | 97.92 |
| Naïve CD8+ | 67 | 67 | 100.00 |
| Memory CD8+ CD57− | 46 | 50 | 92.00 |
| Memory CD8+ CD57+ | 55 | 57 | 96.49 |

Recall is the number correctly classified divided by the test-cell count for that reference class. Six of the eight four-class errors occurred between memory subsets: four CD57− cells were predicted as CD57+ and two CD57+ cells as CD57−. The dedicated binary tests contained 33 naïve and 54 memory CD8+ cells, or 25 CD57− and 29 CD57+ memory CD8+ cells, respectively.

#### Supplementary Table 5. Mixed-sample subset fractions by day

| Day | Subset | FACS (%) | HT + AI (%) | Difference (pp) |
|---|---|---|---|---|
| 1 | CD4+ | 72.7 | 99.0 | +26.3 |
| 1 | Naïve CD8+ | 13.8 | 0.6 | −13.2 |
| 1 | Memory CD8+ CD57− | 6.9 | 0.1 | −6.8 |
| 1 | Memory CD8+ CD57+ | 2.4 | 0.0 | −2.4 |
| 2 | CD4+ | 41.6 | 51.0 | +9.4 |
| 2 | Naïve CD8+ | 39.0 | 36.8 | −2.2 |
| 2 | Memory CD8+ CD57− | 9.2 | 5.1 | −4.1 |
| 2 | Memory CD8+ CD57+ | 10.2 | 6.3 | −3.9 |
| 3 | CD4+ | 41.6 | 40.0 | −1.6 |
| 3 | Naïve CD8+ | 39.0 | 48.8 | +9.8 |
| 3 | Memory CD8+ CD57− | 9.2 | 3.0 | −6.2 |
| 3 | Memory CD8+ CD57+ | 10.2 | 8.0 | −2.2 |

FACS, fluorescence-activated cell sorting; HT + AI, holotomography combined with artificial intelligence classification; pp, percentage points. Differences are HT + AI minus FACS. Each day corresponds to one donor. Values are reproduced without renormalization; the April 25, 2026 summary reports identical FACS values for Days 2 and 3. Day 1 contributed to model development with assumed CD4+ labels. Development–evaluation overlap, fraction denominators, and inclusion criteria were not available (Supplementary Methods S5).
